# Tracking bacterial lineages in complex and dynamic environments with applications to growth control and persistence

**DOI:** 10.1101/2020.03.27.006403

**Authors:** Somenath Bakshi, Emanuele Leoncini, Charles Baker, Silvia J. Cañas-Duarte, Burak Okumus, Johan Paulsson

**Affiliations:** Department of Systems Biology, Harvard Medical School, Boston, Massachusetts, USA; Department of Engineering, Cambridge University, Cambridge, UK; Biophysics Program, Harvard University, Boston, Massachusetts, USA; ArcherDX, Saint Louis, Missouri, USA

## Abstract

As bacteria transition from exponential to stationary phase they change greatly in size, morphology, growth and expression-profiles. These responses also vary between individual cells, but it has proven difficult to track cell lineages along the growth curve to determine the progression of events or correlations between how individual cells enter and exit dormancy. We developed a platform for tracking >10^5^ parallel cell lineages in dense and changing cultures, independently validating that the imaged cells closely track batch populations. Initial applications show that for both *Escherichia coli* and *Bacillus subtilis*, growth changes from an ‘adder’ mode in exponential phase to mixed ‘adders-timers’ entering stationary phase, and then a near-perfect ‘sizer’ upon exit – creating broadly distributed cell sizes in stationary phase but rapidly returning to narrowly distributed sizes upon exit. Furthermore, cells that undergo more divisions entering stationary phase suffer reduced survival after long periods of dormancy but are the only cells observed that persist against antibiotics.

## Main

Bacteria in natural environments experience periods of starvation and stress, punctuated by the arrival of new nutrients that then are depleted again as cells grow and divide^1,2^. Many mechanisms have therefore evolved to help bacteria weather the busts and exploit the booms^3–5^. Traditionally these have been studied by inoculating cultures of stationary phase cells into fresh media and following them along the growth curve back into stationary phase^6–8^. However, batch assays consider average properties of cells, while single cell studies have revealed great cell-to-cell heterogeneity during stress^9,10^. This could be a side effect of saturated pathways displaying less heterogeneity than sub-saturated pathways, making all cells ‘happy the same way but unhappy in different ways’, but could also be an adaptive response to maximize inclusive fitness in uncertain times^11^, e.g. by making a small fraction of cells persistent to drugs^12^. Either way, because the heterogeneity is so substantial and so profoundly changes the fate of stressed cells, effective studies of bacterial responses to stress and starvation should ideally monitor individual cells as they enter and exit stationary phase.

This is challenging for several reasons: *First*, it is straightforward to sample cultures at different times, manually^13^ or with microfluidic automation^14^, but such snap-shots provide little information about the progression of events or the connection between heterogeneity and growth^15^. The dynamics of fluctuations are also important e.g. since rapidly changing heterogeneity is easily time-averaged by affected processes while slowly changing heterogeneity can establish effective cell states. *Second*, studies of heterogeneity are hard to interpret or reproduce unless local conditions are tightly controlled, which is particularly challenging to achieve when components become limiting and conditions keep changing. *Third*, many important outliers in stress response and stationary phase can be exceedingly rare, making it important to sample large numbers of cells. Some microfluidic devices boost throughput by accumulating data over time^16^, but experiments during changing conditions must instead rely on parallelism.

Here we present a microfluidic platform based on the ‘mother machine’ design^17^ that addresses these challenges. We monitor physiology and gene-expression of cell lineages over multiple consecutive rounds in and out of stationary phase, while ensuring that the cells imaged behave quantitatively as the cells in a connected batch culture that can be simultaneously sampled. This provides a microcosm of bulk growth with exceptional resolution and control while enabling conventional bulk assays on the same culture. Our current throughput of >100,000 cell lineages, imaging 10^8^ cells per day, each every few minutes, is high enough that we observe spontaneous persisters without using special mutants^18,19^. The setup also works at very high cell densities, e.g. OD_600_≈10, both for the Gram-negative *Escherichia coli* and Gram-positive *Bacillus subtilis*. In a first application we probed the size-regulation principles during entry into and exit from stationary phase, and demonstrated trade-offs between the behaviors entering and exiting stationary phase.

Platforms like the ‘mother machine’^17^ where individual cells grow and divide in narrow trenches that are fed diffusively by orthogonal flow-channels, can achieve exceptional spatial and temporal uniformity. However, these consider balanced growth in fresh medium, in stark contrast to microbiology experiments more broadly^21–23^. The challenge is that fluid handling for non-viscous and unchanging liquids is very different from the handling of dense and changing cell cultures needed to study stationary phase cells or ecosystems. Also, it is easy to starve cells on any platform by simply letting media run out, but for reproducibility and interpretability, the imaged cells should be starved the same way as in corresponding bulk experiments. We therefore built a system to flow dense cultures from conventional batch experiments into a mother machine type of microfluidic device, (Fig. 1a-b), such that cells loaded in the mother machine will experience the same environment as cells in the batch^24^. Since cells can be retained in the device for hundreds of generations, this allows us to observe how individual cell lineages pass through multiple rounds of entering and exiting stationary phase (Supplementary Movie 1). The process is fully automated, has virtually zero dead volume and provides near-perfect timing about when the media changes occur (SI section 1). The optical density is monitored continuously using a custom OD-meter, in series with the flow path, to align any part of the observed single-cell data to the bulk OD (SI section 2 and 7). This allows us to correlate specific states of nutrient depletion and bulk properties with their corresponding single-cell phenotypes at any time.

**Fig. 1.**
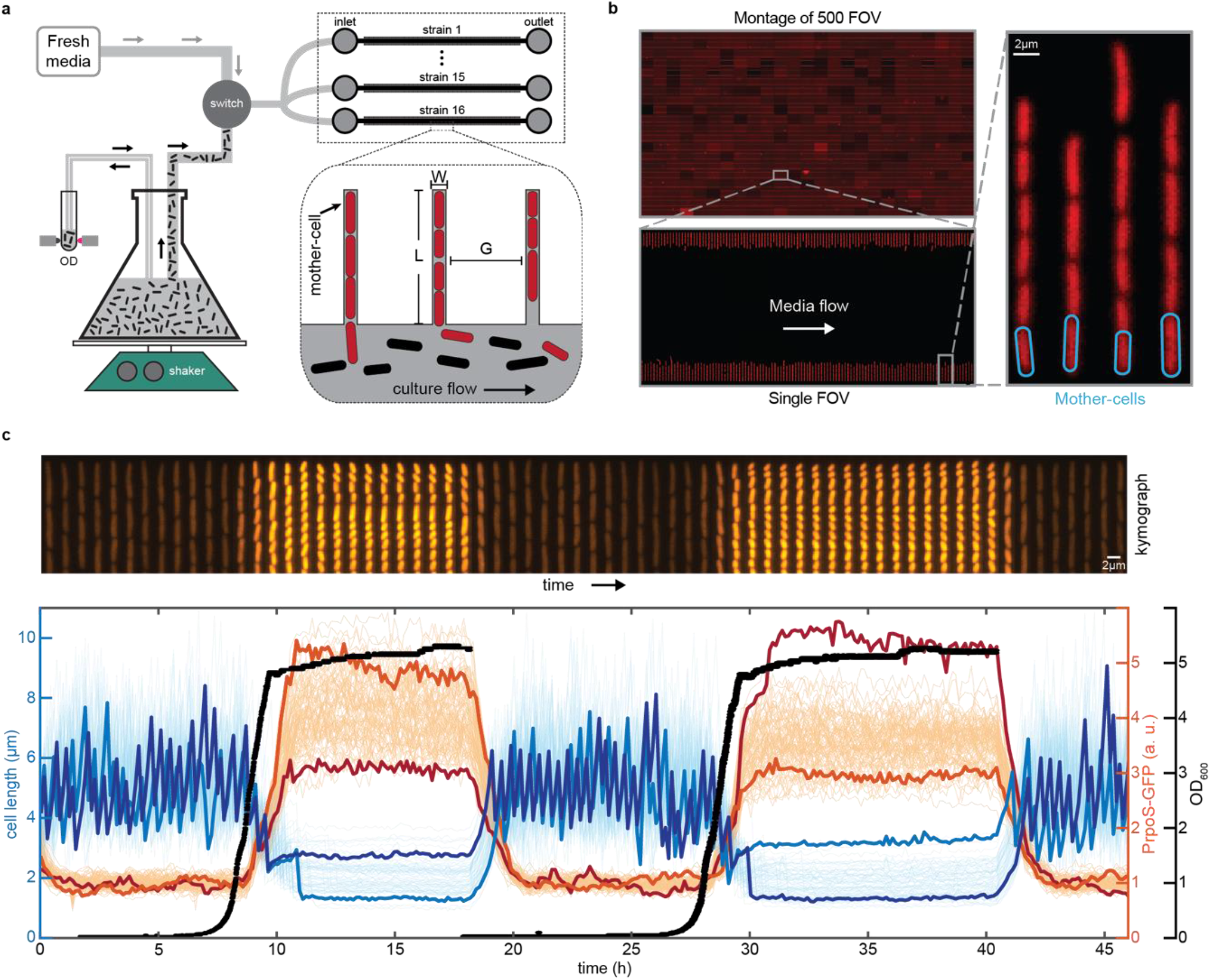
Accurate high-throughput measurement of cell-growth physiology and gene-expression along growth-curve. (a) A simplified schematic depicting the growth curve platform. The platform is based on the mother machine microfluidic device (right of the panel) in which cells under observation are grown in trenches (shown in red) while liquid media is pumped through an orthogonal flow channel. To observe growth dynamics, we flow actively growing bulk culture into the mother machine device while continuously observing its optical density (OD). As the bulk culture flows past (black cells) the red cells in the device (red) respond synchronously with the batch culture. At any point, a switch can be made to flow fresh media, allowing us to observe cells return to optimal exponential growth. The dimensions (W, L, and G in the inset) of the mother machine were highly optimized to meet the demanding requirements associated with flowing dense cultures through the flow channels. (b) The optimizations made to the mother machine design have greatly improved the throughput. We can image up to 16 strains in parallel, with imaging of 705 fields of view (FOV), each containing 186±1 lineages, in under 5 minutes, giving a throughput of 131,072 (= 16 ∙ 8,192) lineages imaged every 5 minutes often for multiple days. We show a montage 500 FOVs in the top left corner. Note the range of intensities present in the individual FOVs in the montage of FOVs has been increased for visualization purposes. (c) (TOP) A kymograph showing a single lineage of cells in a trench expressing a fluorescent RpoS transcriptional reporter as it goes through two consecutive rounds of growth-curve as depicted in the panel below. (BOTTOM) Here we show 80 single-cell traces, from a single field of view, of RpoS expression and cell-size as cells enter and exit from two consecutive rounds of stationary phase. Two cell size traces and two expression traces are highlighted to illustrate high variability between the two rounds of stationary phase. The high-throughput measurements of each property allow us to measure accurate distributions of expression level and cell sizes at any time-point along the growth-curve.

Several technical challenges must be overcome to ensure that cells in the mother machine are representative of the changing flask: First, the shaking batch cultures must be separated from the microscope to prevent vibrations from transmitting (Supplementary Information section 3), and the flow-path from flask to chip must be carefully designed to prevent bio-film formation, non-uniform temperatures, exposure of cells to surfaces other than inert tubing, cross-contamination between the fresh-media and culture paths etc. After extensive trial and error, our setup (SI section 1) addresses those challenges in a way that can be easily adapted for almost any setup. Second, shaking cultures contain bubbles that disrupt flow, requiring custom bubble-traps (SI section 2). Third, maintaining proper diffusive feeding and retention of cells for days is particularly difficult along the growth-curve because cell volumes change greatly during entry and exit. We therefore evaluated 540 combinations of widths, heights, and lengths of the narrow trenches where cells grow and divide, (L, W, and G in Fig. 1a) to find near-optimal parameters for each growth condition (SI section 4). Fourth, since it is not possible to accumulate data over time, as for constant conditions, we systematically optimized the layout to maximize the number of cells imaged while maintaining effective feeding (SI section 6) and ensuring sufficient distances between trenches to avoid significantly overlapping point-spread functions of light between neighboring lineages (SI section 5). The platform records cell size, morphology, and changes in gene-expression levels as cells traverse the growth-curve, while tracking either over 8,000 lineages from each of 16 different strains or over 120,000 lineages of a single strain (SI section 6), imaging every few minutes over the course of multiple days as cells experience multiple rounds of entry and exit from stationary phase (SI section 6). Because each trench contains multiple cells, this allows us to observe on the order of 100 million cell divisions per experiment – enough to observe many rare events and to accurately measure distributions of properties, while also correlating multiple events along the growth-curve, see, Fig. 1 for examples.

Finally, our goal is not merely to subject cells to some unknown level of stress and starvation, but to observe cells along virtually *the same* growth curve as in the connected batch culture, for quantitative comparisons. To confirm this, we therefore first tracked the dynamics of cell-size and a transcriptional reporter for the master regulator of stress response RpoS, Prpos-GFP, upon entry to stationary phase. As the OD of the batch culture goes through a diauxic shift, the bulk growth-rate drops momentarily, and both the cell length and Prpos reporter fluorescence respond synchronously (Fig. 2a, Supplementary Movie 2). Indeed all observed bulk changes appear to impact cells on chip as expected and with minimal delay (SI section 8). To ensure quantitatively similarity also at the level of single cells, we compared cells on chip to cells sampled from batch for immediate microscopy. Cell length and Prpos-GFP reporter activity at different time-points along the growth-curve (Fig. 2b) matched closely (Fig. 2c), to the point where the slight discrepancy of PrpoS dynamics reflected an imaging artefact in agar-pads (SI section 5 and 12) where some of the light emitted by a cell is allocated to its neighbors – another reason to use the microfluidic setup instead. For exit from stationary phase, high statistical sampling is difficult with traditional microscopy since the cultures are so dilute. We therefore used a second microfluidic device which makes it possible to take snap-shots of cells from batch cultures in very high throughput^28^. The results again closely matched each other (Fig. 2d), showing that our platform quantitatively captures the single cell dynamics of conventional batch culture.

**Fig. 2.**
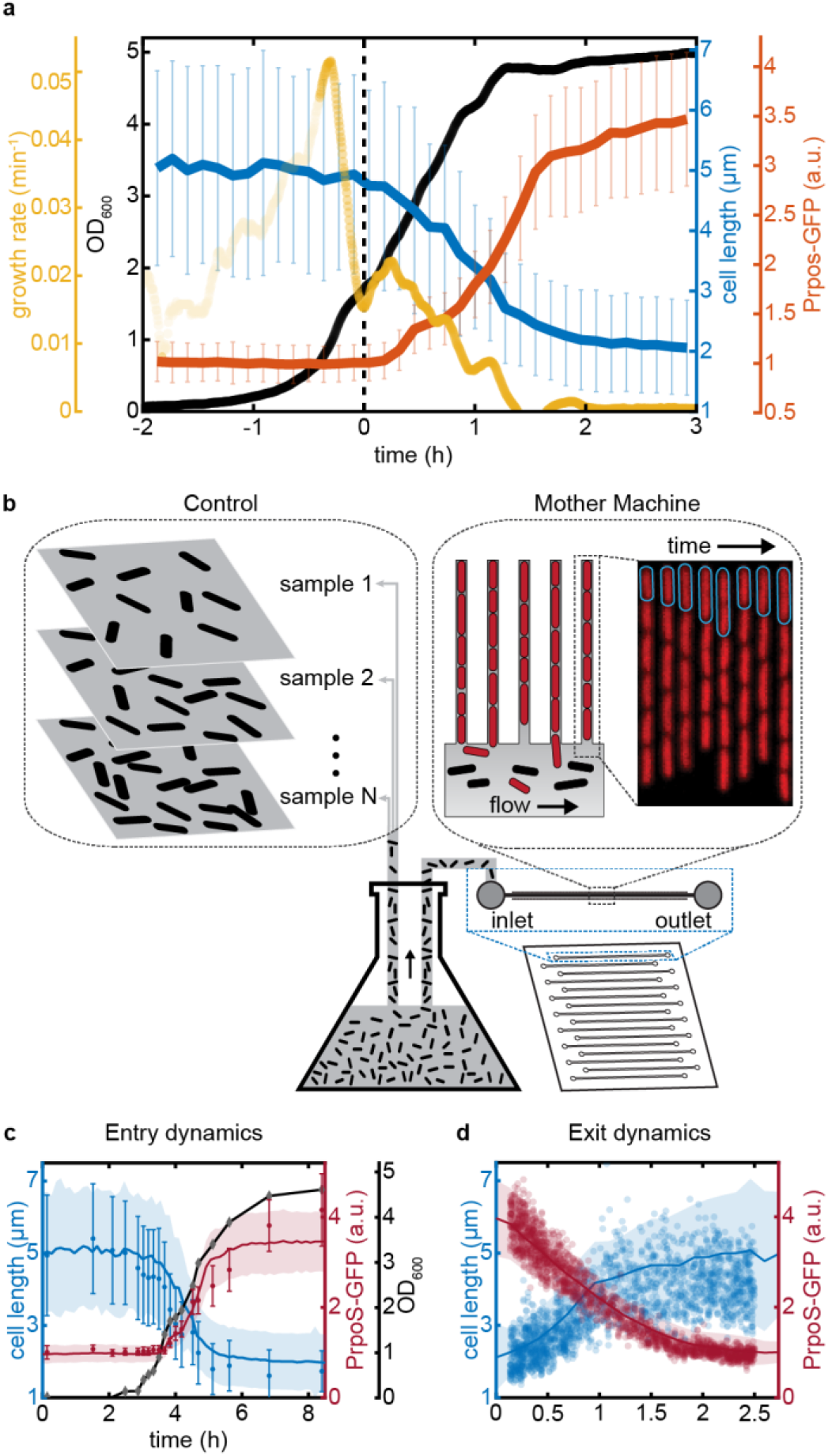
(a) The bulk dynamics observed in the flask are synced with the dynamics of single cells in the mother machine. When the optical density (black) of the culture goes through an inflection point – valley in bulk growth rate (yellow), representing here the rate of change in log(OD) per minute – there is a synchronous drop in the average cell length (blue) and an increase of RpoS transcriptional activity (orange) of the cells in the mother machine device. (b) To examine how well the cells in the mother machine trenches mimic the cells grown in the bulk culture, we compared the average dynamics of cells in mother machine with snapshots from cells grown in the growth-chamber flask. The average trend from the cells in mother-machine are plotted as solid lines (blue – cell length, red – RpoS transcriptional activity). In (c) simple agar pad snapshots (blue circles – cell length, red circle – RpoS) are compared with measurements in mother-machine during the entry to stationary phase. OD600 (black) of the culture is shown in [0, 7.2] range as in both y-axis. During exit, the snapshots were acquired with the MACS microfluidic device^28^, which allows very dilute cultures present during exit from stationary phase to be observed (d). The value from individual cells are plotted as blue circles (length) and orange diamonds (RpoS). The shadowed region and the errorbars corresponds to +/− 1 standard deviation. The average cell length from snapshots of bulk culture is generally shorter than average lengths in mother-machine, due to different age distribution, with more newborn in bulk culture compared to cells in mother-machine.

The ability to quantitatively track growth and expression dynamics with single-cell resolution as bacteria enter and exit dormancy could provide important information about virtually any process in the cell. We first applied it to study how cells modify their size along the growth-curve, as they enter stationary phase from exponential phase and exit upon the introduction of fresh media. Specifically, previous analyses have characterized where cell size regulation falls with respect to three simple phenotypes: adders, sizers and timers, that capture how individual cells in a population respond to being born smaller or larger than average. The adder mode is characterized by adding a constant cell volume each division (in the sense of being independent of birth size) the sizer mode is characterized by division at a fixed volume threshold, and timers are characterized by division after a fixed time interval^29^. Virtually all studies on size control to our knowledge have concluded that all bacteria, archaea, and eukaryotes studied rather closely follow adder-like phenotypes in a wide range of environments from fast growth to starvation^30–33^. However, growth phenotypes have to our knowledge not been tested as cells gradually enter into and exit from stationary phase.

We used a 100x apodized phase objective, imaging 2,215 lineages of *E. coli* MG1655 at one-minute intervals for over 30 consecutive hours to observe growth dynamics during balanced growth and then entry and exit from stationary phase. The 100x apodized phase imaging using the oil objective and one-minute sampling rate reduces throughput but provides fine resolution and accuracy for the size-regulation study. We computed the specific growth rate (rate of increasing cell mass), and the splitting rate (inverse of time between cell divisions) (Fig. 3a) as detailed in SI section 13. In balanced growth, the observed average specific growth rate (0.044 per minute) and splitting rate (0.043 per minute) are both similar to the estimated bulk growth rate (0.044 per minute) (SI section 8 and 13). The CV in specific growth rate (0.053) was approximately three times lower than the CV in the splitting rate (0.16), in agreement with previous literature^34,35^. After the period of balanced growth, we observed an initial sharp drop in the specific growth-rate followed by a brief increase (Fig. 3c) before it steadily declined to zero, as expected as cells go through diauxic shift but then deplete all nutrients (Fig. 3b-c, SI section 8). The splitting rate by contrast is refractory through the diauxic shift for several more divisions (Fig. 3c). This causes cells to produce daughters with ever decreasing sizes as cells enter stationary phase, causing the average cell size to drop rapidly (SI section 14). A similar phenomenon was previously observed in bulk experiments during nutrient downshifts, and attributed to ‘rate-maintenance’^36^, caused by the fact that cells can only split into two daughter cells once initiated rounds of replications are completed.

**Fig. 3.**
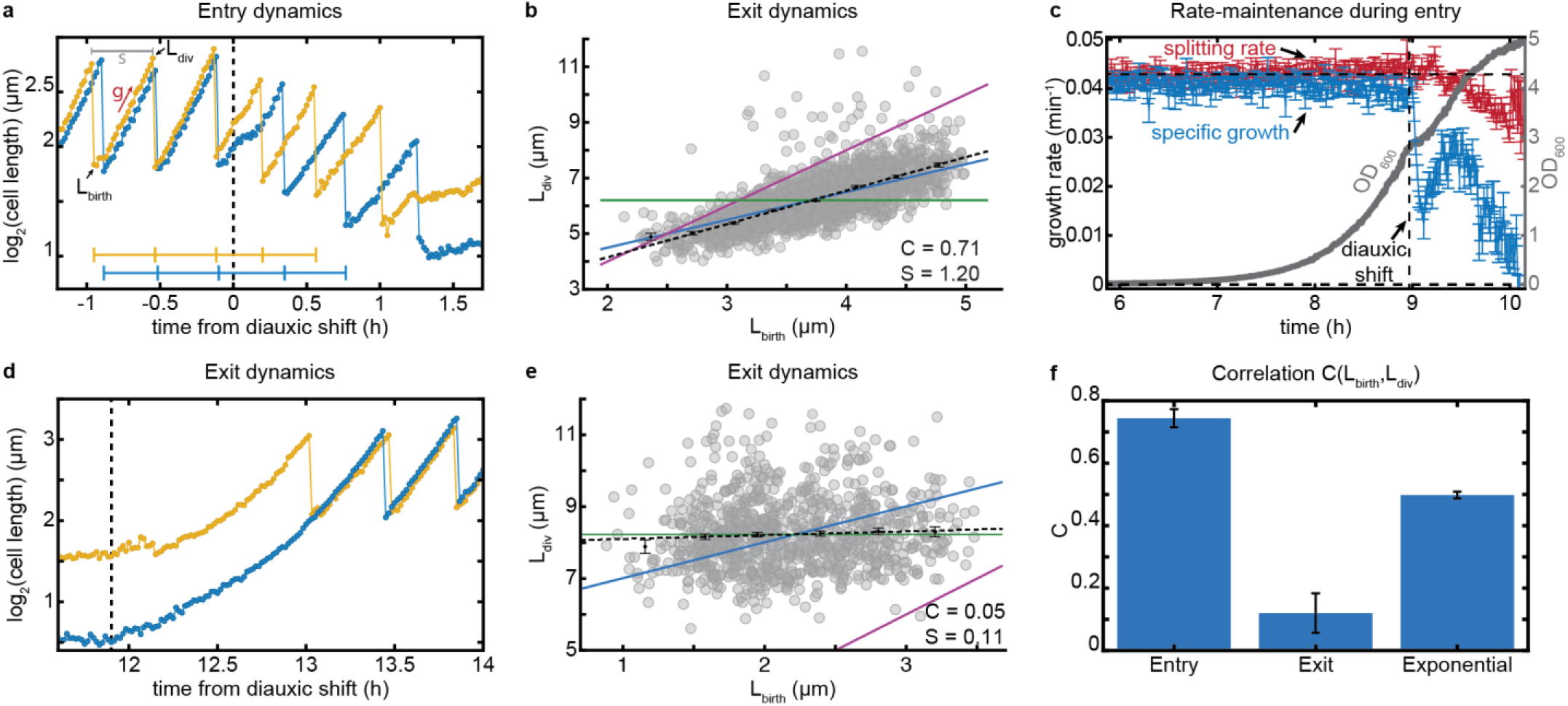
Cell size regulation during entry and exit from stationary phase reveals novel control principles. The timestamps before entering stationary phase are shifted with respect to diauxic shift (t = 0 at d.s.), and after the exit are shifted with respect to the time-point when fresh-media was flowed in. (a) Cell lengths of two representative bacteria are shown (log-scale) as they enter stationary phase after passing through diauxic shift (d.s.). Individual cells slow down their growth during the diauxic shift period. Both the specific growth-rate (g) and splitting-rate (s) can be computed from a single trace (see SI section 13 for details). Since the data is collected every 1 min, we get a high-resolution estimation of the size at birth (L_birth_) and size at division (L_div_). (b) Correlation between cell lengths at birth and division are plotted during the entry to stationary phase (1.0 hour from diauxic shift). (c) The average splitting rate and specific growth-rate of the population are plotted in terms of number of doublings per minute at each time point of the experiment relative to the diauxic shift time-point. During the entry to stationary phase, the cell splitting rate (red) is changing gradually through the diauxic shift (dotted line), but the natural specific growth rate (blue) drops precipitously. (d) Cell lengths from two bacteria are shown during the exit from stationary phase. (e) Correlation between cell lengths at birth and division are plotted during the exit from stationary phase. (f) Size regulation data from 5 different growth-curve experiments are shown. Cells behave as mixed adder-timer during entry to stationary phase (<C_entry_> = 0.74), and act as sizer during exit from stationary phase (<C_exit_> = 0.12) and return to exponential phase where they behave as adders (<C_exponential_> = 0.50).

To investigate the size-regulation strategies employed by cells in different growth phases, we report the Pearson correlation coefficient (C) between cell size at birth (L_birth_) and division (L_div_)^29^. The correlation metric spans a continuum of phenotypes, where the classic modes of timer, sizer, and adder corresponds to three specific values (C = 1, C = 0, C = 0.5). We first confirmed that the size-regulation of our cells during balanced exponential growth was consistent with the “adder” mechanism (C=0.54, Fig. S9A). For non-balanced conditions, the analysis must be done carefully to avoid confounding effects from an average that changes systematically in time. We used the high throughput to avoid such effects by only comparing cells born in the same brief time interval, analyzing their subsequent growth as a function of their varying initial size (SI section 10). Consistently with the rate-maintenance phenomenon, where cells maintained a constant division time even as they start to grow more slowly, this revealed that the size control phenotype looks like a mix of an adder and timer (C=0.71, Fig. 3b). During exit from stationarity the cells started out with widely varying sizes but still underwent the first division at a uniform size, i.e., with almost pure *sizer* dynamics (C=0.05, Fig. 3e). We further calculated the slopes of the linear fit of division length vs. birth length, observing slopes of 0.998 for exponential phase, 1.20 for entry to stationary phase and 0.11 for exit from stationary, consistent with the proposed adder, mixed adder-timer and sizer modes respectively. We note that while the cells act as near-perfect sizers in the first division during exit, they then immediately switch back to adder mode from the second division onwards. The size regulation strategies observed were also consistent with measurements at different temperatures (30°C and 40°C) (SI Table S2).

To explore whether such size control phenotypes are shared by other bacteria, we performed similar measurements on the Gram-positive model organism *B. subtilis*, which is separated from *E. coli* by a billion years of evolution. Experimental details are provided in Supplementary Information section 17. In remarkable similarity to *E. coli*, the size regulation of *B. subtilis* cells during exponential phase was also consistent with an ‘adder’ mechanism (C=0.52) as previously observed, but a near-perfect sizer (C=0.09) for exit from stationary phase (Fig. S9b and S11). Analysis of the size-regulation during entry to stationary phase for the *B. subtilis* strain is complicated by lysis of cells and consequent growth-resumption of cells in the same trench (Supporting Movie 3), a known ‘cannibalistic’ behavior of starving bacillus strains^37^. However, we found that *B. subtilis* cells also display ‘rate-maintenance’ during the entry to stationary phase, which suggests that they could also follow ‘mixed adder-timer’ dynamics.

It has been challenging to identify the underlying mechanisms of growth control partly because so many mechanisms are consistent with ‘adder’ phenotypes^38,39,40,41^. The distinct pattern of adders, timers and sizers we observe in different growth phases may provide more discriminatory tests. For example, many growth control models invoke replication patterns, which are different in entry and exit from stationary phase^39,40^. Dividing more times without additional protein production also causes changes in the levels of division proteins, which also have been invoked in explanations of growth control^41,48^. Repeating the growth curve experiments with various genetic perturbations may thus help pinpoint the pathways involved.

Sizers, adders and timers are also expected to create very different overall heterogeneity in cell volumes, where pure timers fail to compensate for deviations in size and accumulate great variation, while sizers could accurately correct size deviations and adders are intermediate between the two. Partly as a consequence of the mixed adder-timer growth dynamics during entry to stationary phase, the distribution of cell volumes in stationary phase is therefore much broader than during exponential growth (CV about 0.26 in stationary phase compared to 0.16 in exponential growth). The sizer mechanism at exit from stationary phase then causes a rapid return to narrowly distributed cell sizes (CV=0.15). However, some size variation in stationary phase also reflects the fact that some cells undergo more divisions than others (Fig. 4a). These cells could have an advantage, since simply being more numerous allows for more off-spring to explore the world independently. However, those smaller cells would also exit from stationary phase with less internal resources. We next turn to investigating the advantages and disadvantages.

**Fig. 4.**
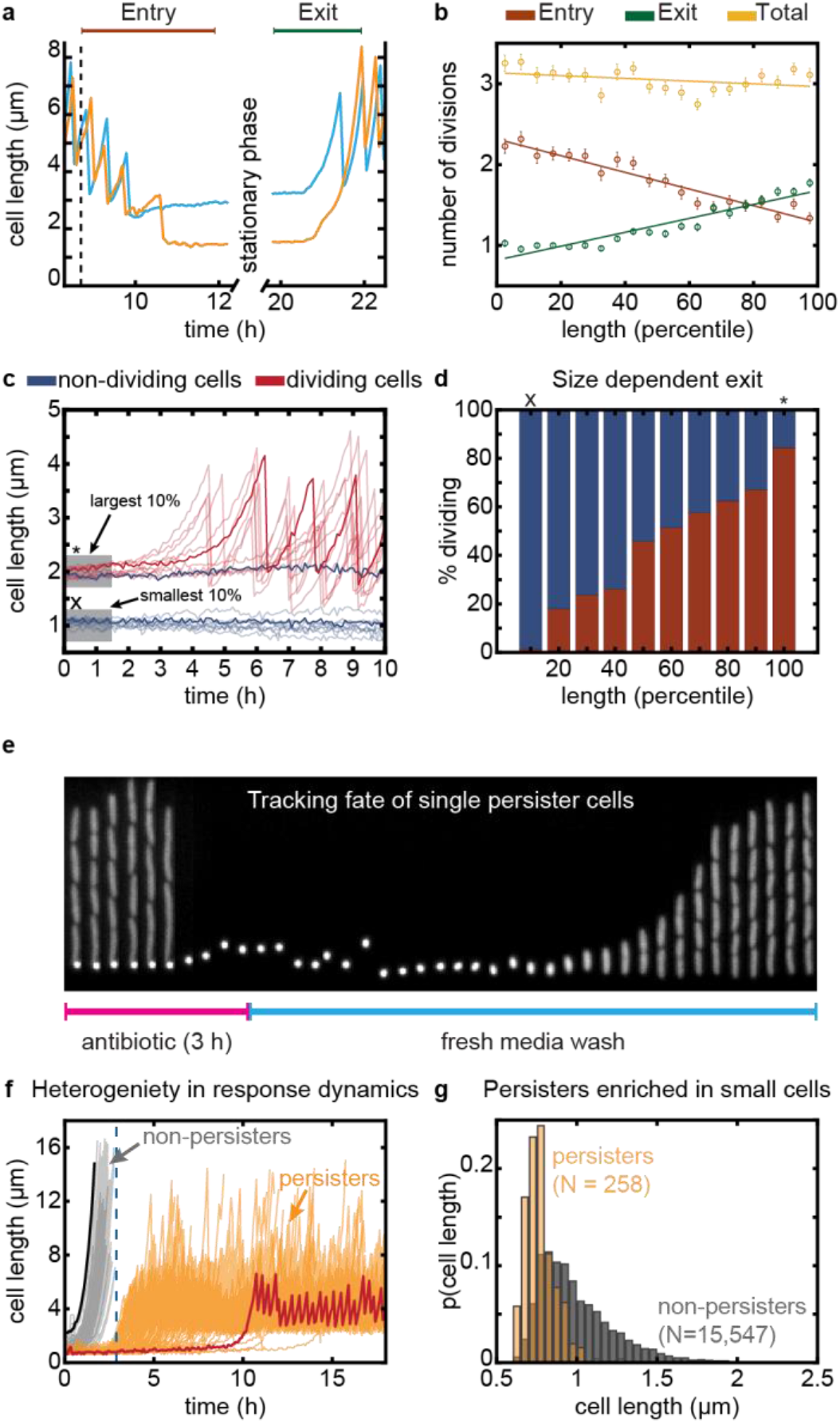
Stationary phase length distribution and survival of deep stationary phase. (a) Two sample traces shown in blue and orange highlight the differing number of divisions performed during entry and exit for large vs. small stationary phase cells respectively. (b) The number of divisions performed during entry (brown), exit (green) and in total from entry through exit (yellow) are plotted vs. stationary phase cell size binned by percentile. The mean values are plotted with error bars (SEM) and a linear regression line is fit to these points. (c) Cells were subjected to a stationary phase lasting 1 week before being provided with fresh media. Cell length time traces for 10 example cells that exited from deep stationary phase are plotted in translucent red, a sample trace is overlaid in opaque red. The same is shown in blue for a sample of 10 cells that did not begin dividing within 10 hours of receiving fresh media. (d) The percent of cells for a given cell size that began dividing within 10 hours is shown in red and the percent that never began dividing are shown in blue. (e) Kymograph showing a trench containing persister mother cell, during antibiotic treatment and recovery. (f) 200 sample traces of persister cells (orange) and non-persisters (cells that chain and die during Amp treatment, in grey) are shown. The largest and smallest cell in the population are highlighted in black and red. Larger cells tend to exit quickly and are therefore vulnerable to Amp treatment, whereas smaller ones exit stationary phase later and survive. (g) Distribution of cell sizes in stationary phase for cells that eventually become persisters in stationary phase is compared to the overall population, where each population has been independently normalized.

We found that for an overnight in stationary phase, cells display great heterogeneity in division patterns both when entering and exiting, but in a negatively correlated manner that cancels out on the net, as to create virtually no variation in the total number of divisions (Fig. 4b). For example, during a time window around stationary phase, where every cell had at least one division before entry and after exit (Fig. 4a), the largest 10% of cells in stationary phase, that divided the least entering stationary phase, on average go through 3.14 ± 0.07 doublings during that time window, whereas the smallest 10%, which divide the most as they enter stationary phase, similarly double 3.26 ± 0.05 times. The same principle was found under all conditions tested, and demonstrates the importance of tracking cells through stationary phase.

We then considered the same question for longer periods of stationary phase. After one week of starvation, we tracked exit from stationary phase upon addition of a poorer defined medium (MOPS buffered medium, MBM)^25^. This by contrast revealed a great net growth disadvantage of cells that divide more times as they entered in stationary phase, where e.g. almost none (<2%) of the smallest 10% of cells began growth within 10 hours of adding fresh media, but 85% of the largest 10% of cells started to divide in that time window (Fig. 4c-d). Larger cells, which spend less resources on increasing their numbers in the entry to stationary phase, can thus reap a substantial net growth advantages after long periods of dormancy (Fig. 4d), perhaps explaining why not every cell divides maximally in the entry to starvation.

That in turn raises the question of why not all cells stop dividing earlier in stationary phase. One answer may be the simple advantage to being more numerous when dispersing to a range of natural environments, but we also considered if smaller cells could better persist when waking up in antibiotic environments^42^. Specifically, a small fraction of cells trigger persistence after starvation^19^, but this has been hard to monitor partly because most platforms cannot ensure local uniformity under stress and partly because persistence is rare. Our combination of tightly controlled local conditions and extreme imaging throughput enables us to directly track large numbers of natural persisters, without mutants that may change both the frequency and properties^18^ of persisters. Because cells in our setup occupy dedicated growth trenches, cells at the end of each trench are also shielded from competition, which enables long-time monitoring of slow growers that would otherwise be lost by dilution in bulk cultures. However, since the cells at the end of the trench have old poles and remained from the initial loading, it is possible that these conditions might influence the observations. To avoid this, we loaded cells that were grown into stationary phase in the bulk culture and treated them with media containing antibiotics, and finally monitored the recovery of persister cells with freshly prepared growth-medium.

Using this approach, we tracked ~80,000 individual lineages that were grown in EZRDM and kept in stationary phase for 36 hours. The cells were then treated for 3 hours with fresh EZRDM media containing a lethal dose of ampicillin (100 μg/ml, 10x MIC, 99.7% mortality), an antibiotic known to specifically target actively dividing cells^43^. To determine which cells return to an actively growing state, antibiotic-free rich growth medium (EZRDM) was then flowed past the cells 3 hours post antibiotic treatment for an additional 12 hours. We found that a majority of cells exiting stationary phase during these 3 hours period was killed by the antibiotic (grey traces in Fig. 4f) treatment, but that some persister cells remained dormant through the antibiotic treatment, and then switched into a non-persister growing state after the antibiotic was removed (see example in Fig. 4e, Supplementary Movie 4). We observed 258 such persister cells, at a frequency of 3×10^−3^, (orange, Fig. 4f), consistent with our bulk measurements (SI section 16) and previous reports^42^.

Comparing cells that did or did not turn into persisters showed a striking trend: the cells that were smaller as they entered stationary phase were much more likely to turn into persisters (Fig. 4g). Indeed, of the 10% smallest cells, 3% turned into persisters, while none of the 45,000 cells that were larger than the average cell produced a single persister. Thus we observe a clear tradeoff: prolonged stays in stationary phase confer great advantages to cells that stop dividing early if cells exit stationary phase without drugs, but instead a great disadvantage if cells exit stationary phase with drugs. This also provides what may be an important new clue in the search for persister-creating mechanisms, by showing that whatever mechanism triggers persistence it is tied to the process of dividing more times during entry into stationary phase, which we suspect is in turn triggered by the need to finish replication forks and place chromosomes in separate cells. That process could deplete resources, but persisters could also be created by the growth and division process itself, e.g. due to fluctuations to low concentrations that arise when cells sub-divide some already low protein abundance into multiple small cells, which does not necessarily change the average concentration but should create more heterogeneity between cells. Indeed, we observe that only some of the small cells become persisters. That could constitute an elegantly simple mechanism for sensing that resources are running out, and using that to create relevant heterogeneity.

The success of microbes depends on their ability to exploit booms and weather busts^44^, but sensing and adapting to environmental changes rapidly enough can be challenging. On one hand cells may not survive the sudden arrival of drugs unless they are already in a drug-tolerant dormant state, and on the other hand they may be unable to take advantage of fresh resources before these are depleted unless they are immediately ready to grow and divide. Clonal populations could therefore benefit from hedging their bets and ensuring that individual cells are prepared for different future environments^45,46^. Great phenotypic heterogeneity has indeed been observed both as cells enter and exit stationary phase^21,47^. However, both the temporal patterns of heterogeneity and their consequences for survival and propagation have been hard to analyze due to the limitations of existing methods^15^. The platform we present here enables us to track lineages entering and exiting stationary phase with precise enough control of local environments to make the results straightforwardly interpretable, over long enough time window, and with high enough throughput to observe key outliers. Because the cell culture can contain virtually any mix of cells, our method could equally be used to study individual cells in ecosystems, e.g. microbiome studies. We believe the ability to track the growth, death and expression patterns of individual cells over time in dense, complex and changing cultures – potentially containing whole ecosystems – could be transformative in many areas of microbiology.

## AUTHOR CONTRIBUTION

SB, EL, and JP designed the study. SB, EL, CJB designed and developed the platform. SB, SJC, and EL designed the microfluidic devices and SJC and EL constructed the devices. SB, EL, and SJC performed the mother-machine experiments. CJB and BO helped with the MACS experiments. SB and CJB developed the image analysis platform. SB, CJB, EL, and JP wrote the paper.

## ACKNOWLEDGMENT

This work was supported by NIH grant R01 AI141966 and DARPA agreement HR0011-16-2-0049. We thank Suyang Wan for her Python code for extracting TIFF files from the ND2 files. We thank Nathan Lord for the master of the first mother machine we used. We thank Po-Yi and Ariel Amir for the discussion of the size regulation analysis. Rich Losick’s lab and David Rudner’s lab provided the strains and necessary protocols for the *B. subtilis* experiments. We thank Pavel Gorelik and Ofer Mazor at the HMS Research Instrumentation Core Facility for assistance in instrument design and fabrication. The microfabrication involved in this work was performed in part at the Harvard Medical School Microfluidics Facility and Center for Nanoscale Systems (CNS), a member of the National Nanotechnology Coordinated Infrastructure Network (NNCI), which is supported by the National Science Foundation under NSF award no. 1541959. CNS is part of Harvard University.

## CODE AVAILABILITY

The code required to process the images (cell segmentation and measurements) are available at https://github.com/manukke/MMsegmenter

## DATA AVAILABILITY

Data can be accessed via the Source Data files provided with this paper.

## ONLINE METHODS

### Strain construction

Strain BO37 was used in the bulk culture flask. It was built by P1 transducing *glmS::*P_RNAI_-mCherry_1-11_-mKate-T1 terminator-FRT Kan FRT::pstS allele into MG1655 as described in Okumus *et al.*^28^. For all the strains used in the mother machine, we used the MG1655 7740 background with ΔmotA. The strain SB8 was used for the phase contrast imaging experiments depicted in Fig. 3b-g and Fig. 4a-b is the background strain for all the other strains containing transcription reporters used in this study. SB8 was constructed by a P1 transduction from the Keio collection strain CGSC#:9565, ΔmotA743::kan into MG1655 and flippase was used to remove the kanamycin resistance^50^. For fluorescent segmentation marker, we made a fluorescent version called SB7, by P1 transducing *glmS::*P_RNAI_-mCherry_1-11_-mKate-T1 terminator-FRT Kan FRT::pstS into this SB8 strain. We made 16 different strains containing transcription reporters for different processes, including the PrpoS reporter, by integrating the transcription reporters constructed by Uri Alon’s group^51^ into the SB7 strain. To rigorously check if the ΔmotA was introducing any artifacts to our measurements, we compared our results with ΔfliC, which also renders the cell nonmotile, and finally compared with the MG1655 6300, which is much less motile than 7740 version and found that the results were identical.

### Mother machine chip preparation

The construction of the wafer and selecting dimension of the single-cell trenches are described in detail in the Supplementary Information. Here we give a brief description of how we made the mother machine devices once we have the wafer ready. To prepare the PDMS for the mother machine chips, dimethyl siloxane monomer (Sylgard 184) was mixed in a 10:1 ratio with curing agent, defoamed and then poured onto the silicon wafer. This was then degassed for 1h using vacuum, and then cured at 95 °C for 1 hour. Individual mother machine chips were cut out of the PDMS mold and holes were created for the inlets and outlets using a biopsy puncher (0.75 mm). Coverslips were cleaned by sonication in KOH followed by sonication in DI water and then bonded to the feature side of the mother-machine chips using oxygen plasma treatment (30 seconds at 50 W and O_2_ pressure set to 170 mTorr) followed by incubation at 95 °C for at least 1 hour. Chips were bonded the day before being used in experiments.

### Loading cells in the mother machine chips

There are two types of experiments described in this article and they use two different types of mother machine devices (SI section 6). In general, 16 different strains can be loaded into the 16 different lanes of the device type 1. We used 16 pairs of gel-loading tips for loading those lanes and centrifuge the chips to load the cells into the trenches that are orthogonal to the loading lanes. The high-throughput experiments involve a single strain to be loaded in a mother machine device where 15-lanes are connected via a continuous snake-like feeding channel. In this case, the overnight culture is concentrated 10x before loading into the device. The chip is then spun at 4500 rcf in a benchtop centrifuge for 2 min, using a custom-built holder. The centrifugal force rapidly loads the cells in the narrow trenches. The dense culture from the feeding channel was washed quickly with fresh growth-medium before starting the image acquisition.

### Description of growth curve platform

Cells are grown in a 500 ml baffled bottom flask (ChemGlass) that is being shaken at 220 rpm on an orbital shaker with magnetic clamps (Benchmark Scientific Orbi-Shaker Jr). A set of peristaltic pumps (Langer Instruments) circulates the culture to and back from a custom-designed inline OD-meter (SI section 2). Another set of pumps takes the culture to the mother machine device loaded on the microscope stage and flows through it. In order to make sure that throughout this entire process cells are maintained at 37°C, to avoid any cold or heat shock, we house the entire systems of pumps, valves, and the shaker with culture flask in a home-built incubator. The incubator was constructed using T-slot framing and acrylic, then heated with an OEM resistive heater and fan. The temperature of the incubator is constantly monitored in two places using a custom microcontroller solution and a data-logging thermometer. This incubator houses multiple peristaltic pumps, solenoid valves and a culture shaker. The tubing containing the culture and fresh media go out of this incubator into the incubator that houses the microscope. The path between the two incubators is kept insulated using an insulating duct the inside of which is actively kept warm using fans that blows the 37°C air from the incubator.

### Description of a growth-curve experiment

Before any experiment is run with the platform, an automated system is used to wash the entire media and culture tubing path. The wash consists of ~40 minutes wash with 20% bleach, followed by 40 minutes wash with 20% ethanol and finally a 40 minutes wash with dH2O. Then, just prior to running the experiment, a bottle containing EZ-rich defined medium (EZRDM) is hooked up to the platform and pumped through all the tubing to remove the water and replace it with media. While this is taking place, the chip is loaded as described above. Once the fresh media has washed out the water, the media used for wash is removed and the flask containing the growth media is attached to the setup.

At the beginning of an experiment, the 500 ml baffled flask containing 200 ml of pre-warmed media is inoculated 1:10,000 of liquid culture. It is then immediately placed into the incubator where it is both maintained at 37°C and actively shaken at 220 rpm. The liquid from a flask containing either fresh media or the actively growing culture is pumped through tubing then into the microscope incubator where it is connected to the mother machine via blunt-end needles. In this path, there is a small, sealed chamber that acts to remove bubbles and is also a vessel in which the OD is continuously measured. A different set of flask and pumps are used to flow fresh media when we need to monitor the cells that exit from stationary phase. The switch between the culture and the fresh media is handled with pinch-valves setup that minimizes any delay between the two conditions and also avoids any possible contamination in the fresh media from the culture (SI section 1).

### Microscopy and image acquisition

We have custom-designed a microscope for the fast acquisition needed for the high-throughput imaging of mother machine. We replaced the turrets that house the dichroic mirror and emission filter to have a single quad band dichroic mirror for all the excitation and have a fast emission filter wheel for all the emission filters. This avoids the lags from the slow movement of the turret and improved our speed of acquisition tremendously. We then used an air objective with high NA (Apochromat, 40x, 0.95 NA) to acquire the images, which made the entire time for stage movement comparable to exposure time, allowing us to scan the entire device in less than 5 minutes. Images were acquired with a sCMOS camera (ANDOR Zyla 4.2), which allowed us to have fast frame-rate with very high QE (84%). We used a LED lamp as the white-light source for the phase contrast imaging and the Spectra-X light engine LED excitation source (Lumencor) for fluorescence imaging. Using these settings we acquired a total of 705 field of views every 5 min, allowing us to track cells in 131,072 trenches over time. For phase contrast imaging, we switched to a high magnification apodized phase contrast objective (100x oil NA 1.3, PH3), that allowed us to get high-contrast images of cells in mother machine. We sampled 85 field of views in 1 min using this oil objective allowing us to image a total of 3,500 trenches. For long acquisition it is important to keep the entire sample in perfect focus, and so the focal drift was controlled via the Nikon Perfect Focus system. The entire setup of flow-switch and the Nikon Ti inverted microscope was housed in a temperature-controlled incubator (OKO lab). We have used the Nikon Elements software for the acquisition control and to create position list using a grid structure. The fluorescent colors were ordered according to their order in the filter wheel, and the travel of the stage along the position list was optimized to get the fastest possible scanning of the entire chip. The PSF was always kept in ‘on’ option for fast scanning. The data were saved locally in ND2 format to improve writing and saving speed and eventually extracted for analysis into single TIFF files using a custom-made ND2extractor Python script.

### Data processing: segmentation and single-cell tracking

We have developed an image-processing pipelines for processing the large volume of high-throughput data collected per experiment (~1.5 TB/ experiment). A multi-core extraction algorithm was used to extract single TIFFs from the ND2 file and organize them into folders for each field of view (FOV). We then proceeded to analyze the stack of single TIFFs for each FOV over time. The pipeline for the analysis involves four major sections: image segmentation, data extraction, single-cell tracking, and trace-cleaning. The image segmentation step involves processing the entire field of view to identify single cells and find a contour for each cell from which the intensity data is going be extracted. One major challenge for single-cell segmentation for a data along the growth-curve is that cells continuously change their size and shape along the growth-curve. To address this, we have developed a custom build segmentation algorithm that is more flexible for variable cell sizes, which is roughly based on the Adjustable Watershed algorithm implemented in FIJI^52,53^. We first adjust contrast across all the frames of the time-lapse image stack to correct for changes in intensity of segmentation marker/phase contrast for every individual the TIFF files. Then we perform background subtraction intensity using a rolling ball algorithm. The background subtracted images were processed with a combination of global and local thresholding to create binary masks for cells from background and use a marker-based watershed algorithm to separate all the binary masks into masks of single cells in a reliable manner. The segmentation algorithms are different for the fluorescent images and phase contrast images. After the segmentation, we extract the fluorescence intensity properties and shape properties for each connected component from the list of masks.

The next step is to track single masks over time to get time-lapse data. An individual FOV has ~230 trenches with ~8 cells each. Therefore, in order to track single cells over time, we have to assign the ~1300 binary masks to their corresponding masks in different time-points. This is a massive task if to be done by the traditional frame-by-frame nearest neighbor assignment. To address this challenge, we have developed a novel clustering algorithm that cluster all the cells in a single trench over the entire duration of the experiment and then assign the top one in each frame as the mother cell. Specifically, we use DBSCAN^54^ clustering implemented in MATLAB^55^ to cluster the data into lineages from individual trenches and then sort the data to get the top cell in each trench and assign all of their properties to mother-cell properties. The segmentation algorithm is not perfect, so we define criteria to help filter out data that is un-physical. The ratio of the division length and birth length should be 0.5, so we filter out any division interval where the length at birth over the length at the preceding division differs from 0.5 by more than 20%, The single-cell traces of the different properties of interest were further cleaned to remove unnatural trends using the reference marker intensity traces.

## Notes

### Competing Interest Statement

The authors have declared no competing interest.

### Summary of Updates

Reduced text and updated figures

