## Supporting information for "Tracking bacterial lineages in complex and dynamic environments with applications to growth control and persistence"

### Contents

| | | |
|---|---|---|
| **Section 1** | **Design of flow paths to prevent biofilm formation and cross-contamination, and to ensure clean recovery from stationary phase** | **3** |
| **Section 2** | **Anti-clogging agents and automated bubble-traps** | **5** |
| **Section 3** | **Dual incubator setup for separating platforms for culture and imaging** | **6** |
| **Section 4** | **Optimization of the mother-machine architecture** | **7** |
| **Section 5** | **Point spread function bleed-through and trench-gap optimization** | **9** |
| **Section 6** | **Mother-machine architecture for high-throughput imaging** | **11** |
| **Section 7** | **Comparison of bulk growth with growth in microfluidic devices** | **13** |
| **Section 8** | **Comparison of growth-dynamics in the bulk culture and at single-cell level** | **14** |
| **Section 9** | ***B. subtitlis* and *E. coli* both follow adder mode during exponential phase** | **16** |
| **Section 10** | **Transition from adder-mode towards timer-mode during entry to stationary phase** | **17** |
| **Section 11** | ***B. subtitlis* behave as sizer during exit from stationary phase** | **18** |
| **Section 12** | **PSF artifacts in agar-pad images** | **19** |
| **Section 13** | **Data quality, analysis, and growth-measurements** | **20** |
| **Section 14** | **Rate-maintenance and cell size in stationary phase** | **22** |
| **Section 15** | **Size regulation in different temperatures** | **23** |
| **Section 16** | **Bulk measurements of persisters** | **24** |
| **Section 17** | **Growth-curve experiments *B. subtilis*** | **25** |

**Section 18 Strain construction control** 26

**Section 19 Media Recipes** 27

**References** 28

### Section 1 Design of flow paths to prevent biofilm formation and cross-contamination, and to ensure clean recovery from stationary phase

We have designed the flow path to ensure that there is no zero-flow pocket for bacteria to get trapped and eventually form biofilms which can not only clog the flow, but also reduce the quality of the nutrient arriving to the microfluidic device in arbitrary ways. We found that bacteria can get trapped into in-line solenoid valves commonly used in microfluidic applications. Therefore, all such solenoid valves were avoided and instead we used pinch valves where cells never have to enter valve compartments and can flow through tubing through the entire path. Not only the choice of valves, but also the connector geometry can have an impact. In particular, connectors with orthogonal geometry can cause particles to get trapped, as shown by Vigolo *et al.* [1]. For this reason, we have avoided the use of orthogonal connectors and we use either 180° zero-dead-volume connectors or 120° Teflon Y-junctions. To ensure proper starvation, we also examined different tubing material, and found that, except Teflon and silicone tubing, all other types of tubing lead to release of tubing material in the flow which the bacteria can consume during nutrient starvation. Therefore, the entire flow path was constructed with Teflon tubing connected with zero dead-volume connectors (made of PEEK material), except the pinch-valves and peristaltic pumps, where we used short segments of silicone tubing that allow pinching.

When the culture medium in the flask is inoculated with cells, the cells naturally experience transition from exponential to stationary phase as the nutrients of the medium runs out. However, in order to bring cells in stationary phase to the exponential phase, when we change the culture path to flow fresh medium, we have to make sure that no cells from the original culture is present in the fluidic path and that there is no delay in the switch (see Figure S1a and b). An uninterrupted recovery from stationary phase to exponential phase ensures robust reversibility of the process, and therefore allows us to follow the same batch of cells through multiple rounds of entry and exit from stationary phase (see Figure S1c and d). At the same time, it is important to eliminate any delay in the switch from culture to fresh medium, to mimic inoculation of cells from stationary phase into fresh medium. To reduce the delay during the switch between the culture flow and the recovery media, we use high flow rate to bring the recovery media close to the microfluidic device and place the pinch-valve switches on the microscope stage so that they are very close the microfluidic chip. The switch has an alternative path that allows us to divert the flow towards a waste during the fast-flow that also moves the culture present in the path to the waste and the flow is only directed to microfluidics device once the entire tubing is free of culture (see Figure S1b).

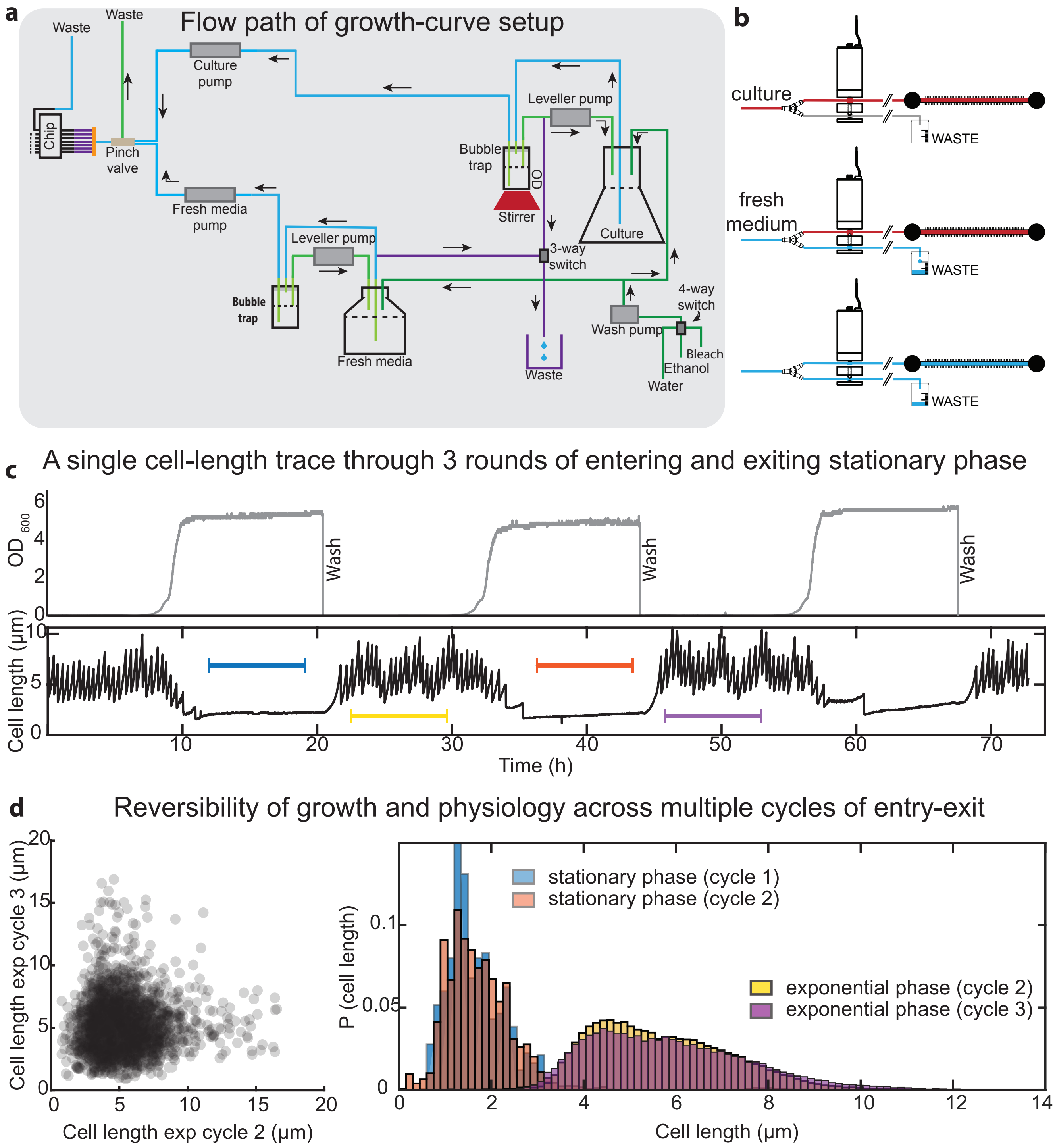


**Figure S1.** (a) Schematic of the flow path for the growth-curve setup. (b) Sequence of flow control for fast switching from culture flask (red) to fresh medium (light blue). (c) Example single cell length data for the same cell entering and exiting stationary phase through 3 rounds of growth-curve. (d) Statistical analysis of reversibility of physiology during multiple rounds of entry and exit. Left: Scatter plot of cell length in exponential phase after the exit from stationary phase in cycle 1 and after exit from stationary phase in cycle 2. Right: Comparing distribution of cell lengths in stationary phase and in exponential phase in different cycles. Color-code in the legend matches the range shown in (c).

### Section 2 Anti-clogging agents and automated bubble-traps

In order to reduce the formation of biofilms or the adherence of cells to the microfluidic device, we use Pluronic F108 (Sigma Aldrich, 542342-250G, 0.8% of a 0.1 g/ml stock) to passivate the PDMS surface [2]. This concentration has been verified to have no impact on cell physiology in exponential growth [3, 4]. In order to ensure that the concentration of Pluronic used in this study does not impact cell growth and physiology through the entire growth-curve, we ran several growth curves in a plate reader using various concentrations of Pluronic F108 (see Figure S2a). We start to observe an impact on growth dynamics and final OD only above concentrations of 3.2% of F108 0.1 g/ml stock. Therefore, we have safely used 0.8% of 0.1 g/ml Pluronic stock solutions. Experiments for B. subtilis were done using 0.1 mg/ml BSA, as described in our previous work [5].

The continuous shaking of the culture causes bubbles to form, which is further exacerbated with the addition of Pluronic F108 or BSA, and can extensively affect measurements in microfluidic devices. The bubbles can also get trapped in the microfluidic device and affect cell growth in the long run. In order to remove these bubbles from the flowing culture before they enter the microfluidic device, we have custom designed automated bubble traps, since the commercial ones will trap the flowing cells in the membrane and thereby will not only cause the nutrient profile to arbitrarily change due to cellular consumption in flow-path, but also membranes to get jammed due to biofilm formation over time. The bubble traps designed in this study can remove bubbles from the culture flow for indefinite amount of time and are also used for a continuous monitoring of the optical density of the culture. A schematic of the bubble trap is shown in Figure S2b.

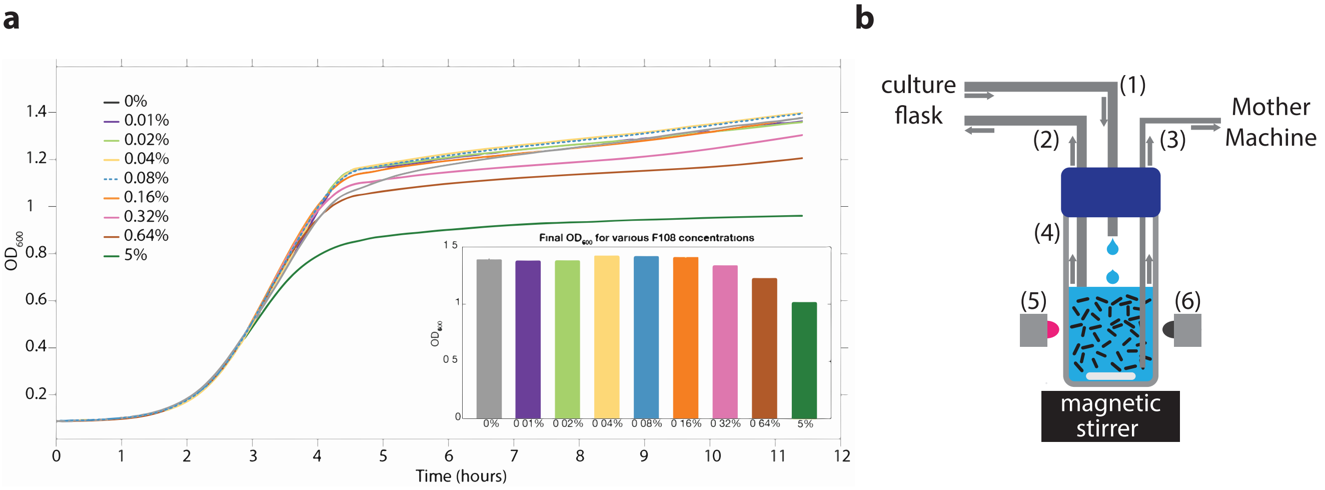


**Figure S2.** (a) $OD_{600}$ vs time plot (average of 12 replicates) for nine different concentrations of F108 is shown. All curves were aligned at $OD = 0.5$ to account for different lag times of growth. (b) Optical density of the growing culture is monitored in real time using a custom-made OD meter: media is pulled from the growing culture at fast speed (1) into the glass vial (4) and it cycles back to the flask via tubing (2). A magnetic bar ensures perfect mixing in the glass vial and the OD measurements are done using transmitted light from the LED emitter (5) to the detector (6). The measurements are read through an Arduino and displayed in real time using a custom MATLAB[6] GUI. The OD meter is also designed to prevent bubbles from flowing to the mother machine chip. In fact, the fast cycling through tubing (1) and (2) ensures that the level of the media is kept constant and that bubbles that get collected on the surface of media are pushed back to the culture flask. The bubble-free flow of the complex media towards the microfluidic chip is ensured by a thinner tubing (3) that is placed lower in the liquid and far from the media-air interface where most of the bubbles collect.

### Section 3 Dual incubator setup for separating platforms for culture and imaging

In order to get reproducible profiles of growth-curves, and for relevance to conventional microbiology assays, we grow our cells in large batch cultures. For optimum growth, large batch cultures must be shaken well to enable thorough mixing and aeration. To isolate the imaging platform from the vibrations arising from the shaker-system, we have engineered a dual incubator system. The culturing platform is housed in one of them (Figure S3, left), while the microscope with the microfluidic platform is housed in another (Figure S3 right). The two incubators are connected via an insulating duct that maintains the temperature around the tubing that carries the cells from the culture flask to the microfluidic device. Hot air from the incubator is blown into the duct through fans at the junctions.

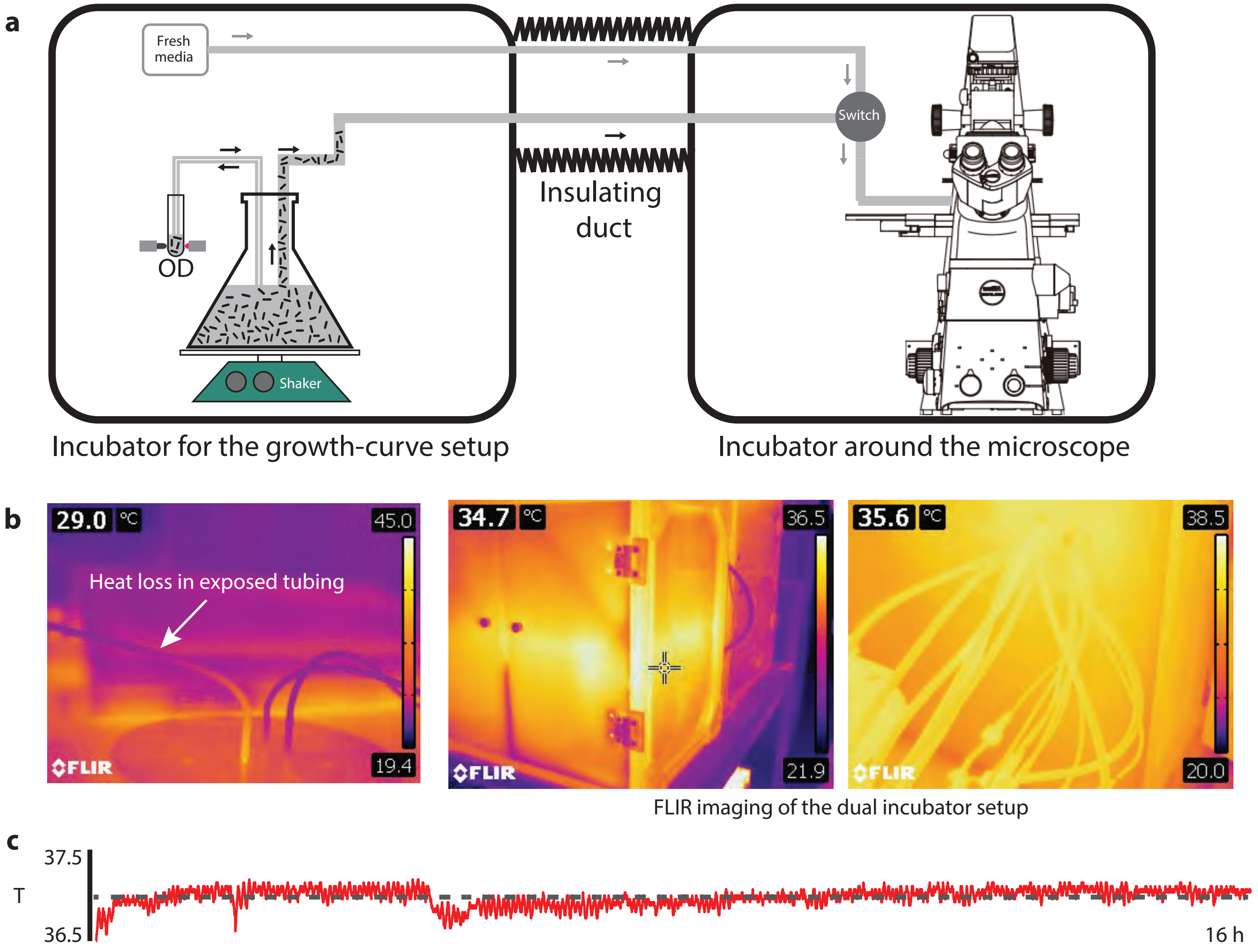


**Figure S3.** (a) Schematic of the dual incubator setup. Incubator 1 (left) houses the batch culture setup (culture flask, shaker, and automated optical density measurement system), and pumps, valves and automated bubble-traps to flow the culture through an insulating duct to the incubator 2 (right) that houses the microscope and microfluidic system. (b) Fluorescent IR imaging was performed to ensure the thermal uniformity of the entire platform. The left panel shows how quickly heat-loss happens through the wall of thin tubes in the absence of insulating ducts. The medium and left panel shows the uniform thermal profile of the custom-designed incubator and tubing. (c) A sample trace of the temperature reading off a probe near the shaker during a regular experiment duration.

### Section 4 Optimization of the mother-machine architecture

The design of the mother machine also had to be highly optimized to handle the vast changes in cell size along the growth curve (15-fold for MG1655), as well as the increased throughput requirements of typical growth-curve experiments and to prevent clogging from dense cultures all while maintaining proper diffusive feeding and retention of cells for experiments that typically last multiple days. In this section, we discuss the optimization of the trench (narrow channels where the cells are loaded) dimensions, and in the later sections, we discuss optimization of the gap between trenches and the layout of the flow-path.

We have sampled approximately 540 combinations of the width, height, and length of the narrow trenches to find the optimal combination of parameters for each growth-condition (Figure S4a, b and c). The length and width of the trenches are optimized for better diffusive feeding and loading, and longer retention of cells in the trenches. If the trenches are not wide or tall enough, then the cells may not load into the trenches and even if they do load properly, the tight fit can affect their growth rate. If these dimensions are too large, then the cells can re-arrange themselves or load side by side in a single trench (see Figure S4e). If trenches are too short, then cells are not retained properly and if they are too long, it can affect the nutrient content at the end of the trench (see Figure S4d). It is thus important to correctly determine these parameters; however, bacteria often vary substantially in size depending on growth media and environmental conditions, thus no single combination of trench dimensions is optimal for all conditions. Along with feeding and retention, the trench-width has to be also optimized for avoiding double loading of cells in each trench, i.e. cells get loaded side by side in a trench, which causes additional stress to the cells and also cause technical challenges for cell-segmentation. This gets even trickier when cells change their width as they transition from exponential to stationary phase along the growth-curve. Along with the trench width and trench length, another important parameter is the height of the trench roof as it plays key-role in loading efficiency and keeping the entire cell in focus, factors which counteract each other. We surveyed about 12 different trench-widths, 9 different trench-lengths for each of those trench-widths, and 5 different trench-heights, and out of 540 combinations of each of those parameters we chose the settings that allowed best growth and long retention times. For the growth curve experiments described in this paper, we chose trench dimensions that are optimal for exponential phase where cells are largest (trench-length = 25 $\mu m$, trench width = 1.5 $\mu m$, trench height = 1.28 $\mu m$). We find that this choice of trench size is just tight enough to properly retain the small stationary phase cells and does not artificially impact growth rates at any point during the growth curve. Using the correct parameters makes it possible to effectively segment each cell throughout the growth-curve as they change their size and aspect ratio.

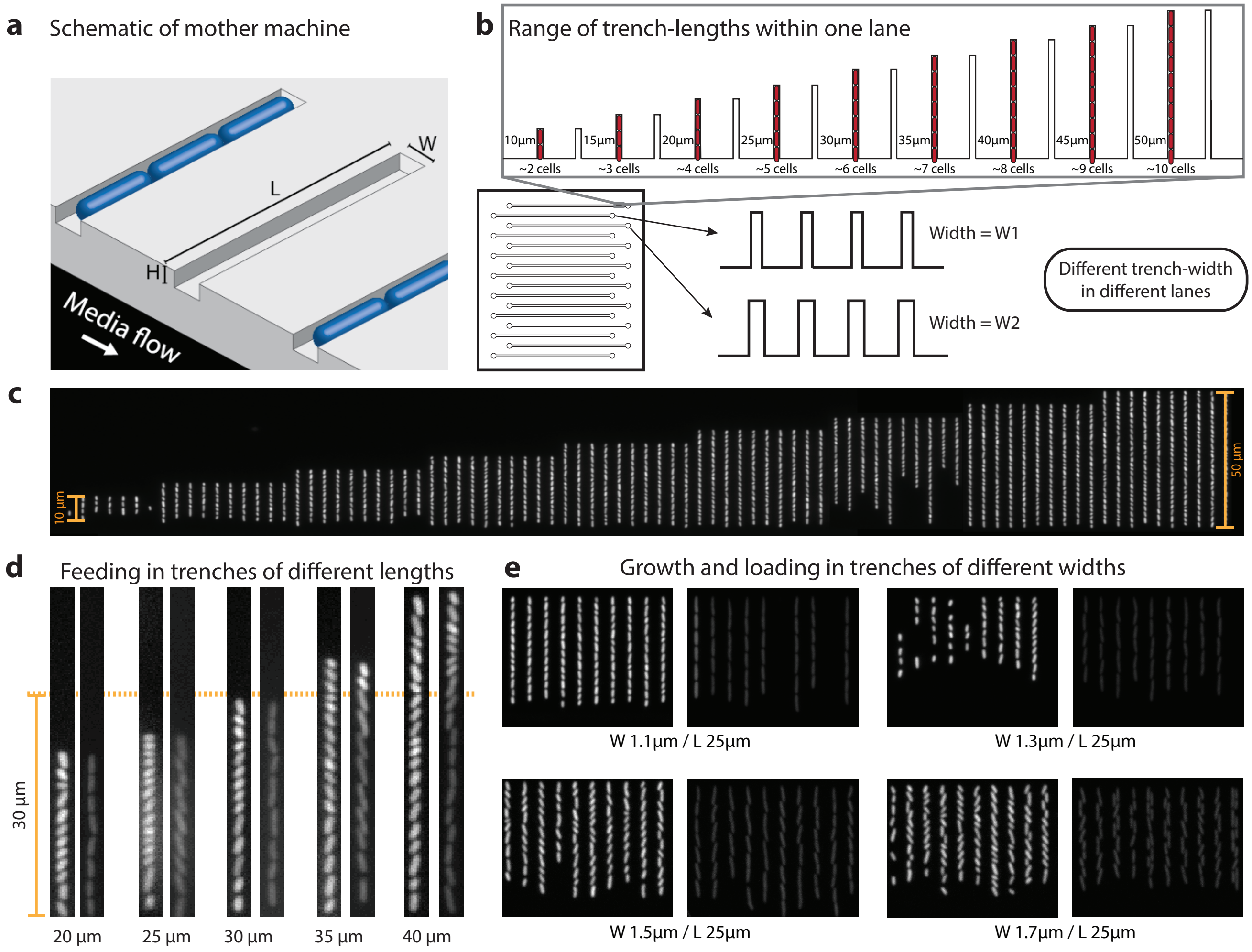


**Figure S4.** (a) 3D schematic to illustrate different dimensions of the mother-machine design. (b) Sampler chip (schematic of the chip in left bottom corner) to verify multiple different parameters in the same experiment. Each lane has arrays of trenches from 9 different trench-length (10 $\mu m$ to 50 $\mu m$). Different lanes have trenches of different widths. (c) Sample field of view of one of the lanes (Note: cells are in stationary phase in this frame, as cells are lost from the short trenches within a short time after fresh media is flown). (d) Nutrient availability depends on trench-length. Five different sample trenches are shown in pairs. The left panel shows cells in stationary phase ($t = 0$) and right panel shows the same trench after 6 hours of flow of conditioned medium. The conditioned medium does not reach beyond 30 $\mu m$ (orange dotted line). (e) Cell loading in trenches of different widths. The left panel shows cells in stationary phase and right is exponential phase. In the narrow trenches (1.1 $\mu m$) cells are constricted and straight (doubling time is slow, 28 min, compared to 22 min in 1.3 and 1.5 $\mu m$ trenches), whereas in the widest trenches (1.7 $\mu m$) the cells tend to double-load.

### Section 5 Point spread function bleed-through and trench-gap optimization

The point-spread function of an imaging system makes the intensity from a point-source emitter spread through a much wider area corresponding to the wavelength of the emission. Therefore, the fluorescent emission from a cell spreads away from the cell creating a decaying signal. This causes fluorescence from nearby cells to be assigned to each other, which distorts the variability of intensity across population. In conventional imaging systems that use microcolonies of cells, like agar-pad, or microfluidic platforms (like commercial CellASIC etc), such bleed-through effects are very significant and unavoidable, which makes the measurements unreliable. Mother machine allows cells to be kept apart from each other in an array of specified distances. However, even then, depending on the distance between neighboring trenches, the bleed-through can be significant. Therefore, we attempted to quantify whether the dimensions of our device allow an artifact-free measurement of such variability.

The cells within a trench of mother machine, the mother-cell and its daughter and grand- daughter, contribute to each other's intensity through such PSF bleed-through. The cells from different trenches can also contribute to each other's intensity through such bleed-through, as a function of the distance between trenches (see Figure S5a and b). The further the trenches are the lower will be the effect of such crosstalk. To quantify the effect of point-spread function bleed-through from neighbor trenches, we have developed an assay that uses signals from YFP expressed from a plasmid that goes to zero when the plasmid is lost. We use an empty position in a trench (trench-of-interest) next to the trench containing the YFP expressing cell (trench-neighbor) to quantify the amount of fluorescence bleed-through by comparing the intensity in the empty trench when the YFP signal is present and lost in the neighboring trench. In order to quantify the bleed-through as a function of the gap between two nearby trenches, we used microfluidic devices with varying gap between the trenches (4 $\mu m$, 6 $\mu m$, 8 $\mu m$, 10 $\mu m$). We find that when the trenches are 4 $\mu m$ apart, the neighbor cell can contribute about 1.1% of intensity to a cell in nearby trench. Since every trench in a mother machine has two nearby trenches on both sides, the total contribution from both trenches is 2.2%. For the microfluidic device where the trenches are 10 $\mu m$ apart, the contribution is much less; 0.2% from two nearby trenches. The most significant contribution to the intensity of a mother-cell, seems to be from the daughter, grand-daughter cells in the same trench, about 2.9%, which can't be avoided as it is limited by the mother machine design (see Figure S5c and d).

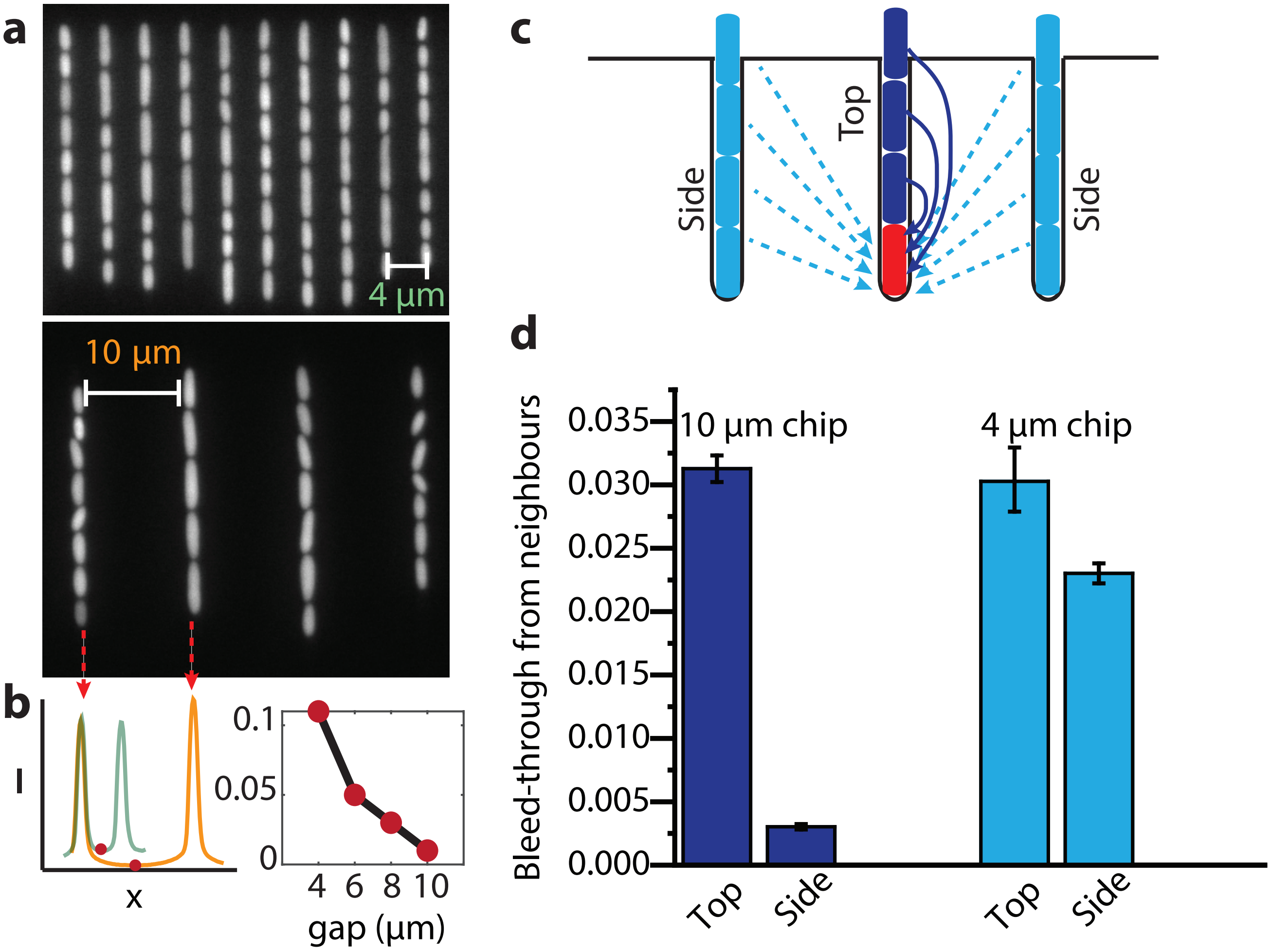


**Figure S5.** The extent of point-spread function bleed-through depends on the distance between trenches. (a) Sample field of views of mother-machine device with different trench gaps (top: 4 $\mu m$; bottom: 10 $\mu m$). (b) Left: line scans of the intensity profiles show that intensity does not go down to baseline in the middle of two trenches. The height of the valley off the baseline gives the extent of the cross-talk (2 times the bleed-through from cells in each trench). The right panel shows how the bleed-through gradually drops with increasing gap between the trenches. (c) Schematic of cells in mother machine showing different component of the bleed-through from different neighbors. Cells in the same trench are referred to as ‘top’, while cells in the trenches from both sides are referred to as ‘side’. (d) Quantification of the extent of bleed-through from the cells from the neighbor trenches ‘side’, and from the same trench for chips with 10 $\mu m$ and 4 $\mu m$ gap, respectively.

### Section 6 Mother-machine architecture for high-throughput imaging

In order to increase the throughput, we started by redesigning the trench layout. First, we placed the trenches on both sides of the flow channel and used a lower magnification objective (40x, NA 0.95) that allows us to image the two rows of trenches in a single field of view (see Figure S6a). We had to reduce the width of the flow channels to bring the two rows closer, and we compensated by increasing its height. The segmentation program was customized to segment cells from the 40x image, and the tracking program was customized to handle both rows of cells in the same FOV. In the standard mother machine device we have 16 different flow channels, in which we can flow 16 different types of media in an independent manner and test 16 different strains in parallel (see Figure S6b). Each of these lanes have 8,192 trenches, giving sufficient throughput for each strain. For experiments that require higher throughput, for example for examining persister cells, we use a layout where we have combined 15 of these lanes in a snake-like architecture (see Figure S6c). The design parameters of the snake was optimized to reduce the flow-time through the entire path and to avoid clogs and blocks at the bends. In this chip we can load a single strain and 122,880 (= 15·8,192 ) lineages imaged in parallel. A single trench has about 5-6 cells and in a single day we get 70 division events, enabling us to observe $10^8$ division events per day. The high-throughput experiments were done using a high NA air objective (NA 0.95, 40x, Plan Apo). However, since fluorescence imaging can cause phototoxic side effects, we could only use fluorescence imaging to sample dynamics at 6 min/frame rate. For faster imaging and higher resolution, we use a 100x apodized phase contrast objective (NA 1.3, 100x oil, PH3). Since this objective has high magnification and therefore smaller field of view, the throughput is much lower than the 40x. The phase objective, being an oil objective, also has lower travel distance, limiting the area of imaging. Therefore, we can image 3,500 lineages per minute using this setup. Phase contrast studies that do not need similar resolution were done using a 40x phase contrast air objective (NA 0.95). This system can image 160 FOVs per minute, enabling simultaneous tracking of around 30,000 separate lineages.

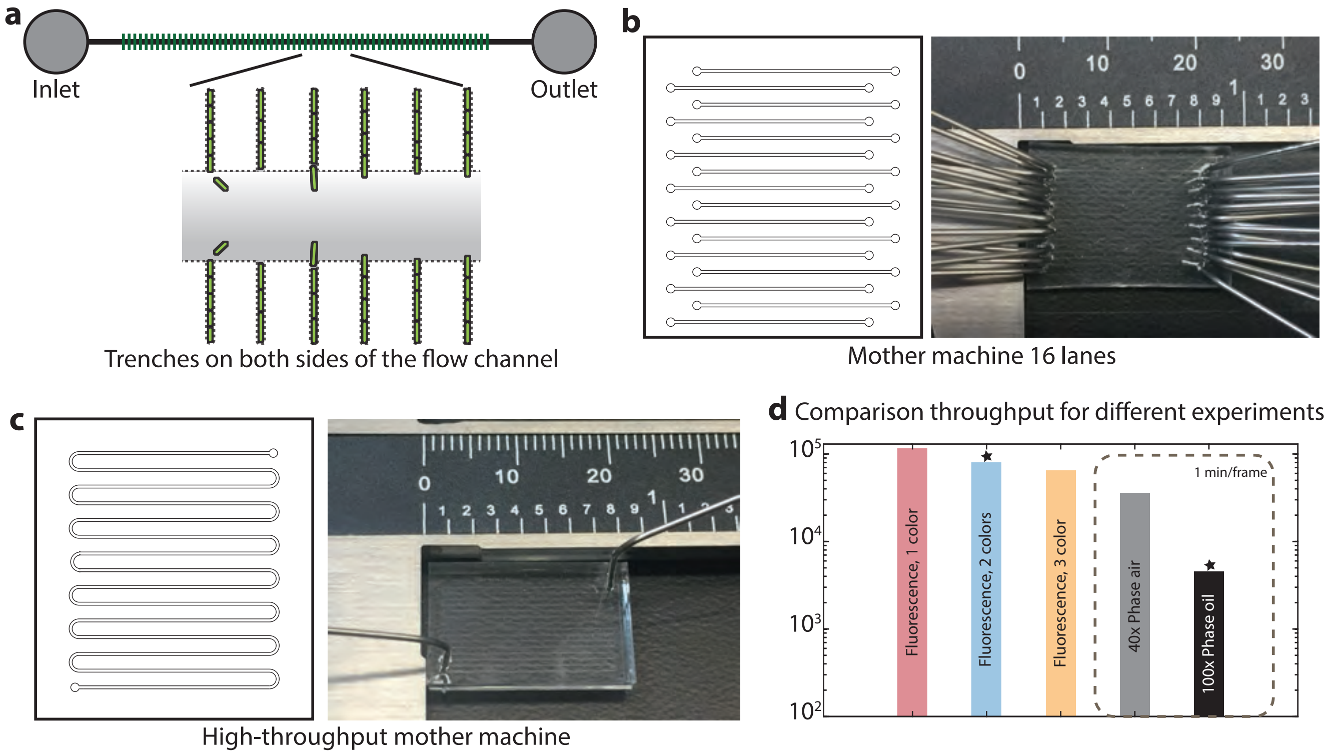


**Figure S6.** Mother machine design: we have designed the flow paths to have trenches on both sides (a), which gives twice the throughput per field of view. Every single field of view in a 40x objective has total $186 \pm 1$ trenches. We have two different designs of the flow path layout through the device. In the type 1 (b), there are 16 total flow-channels allowing us to load 16 different strains to monitor their physiology and gene-expression in parallel. For each strain we get 8,192 trenches to get enough high-throughput for fine-binning of the data. In the type 2 (c), we connect 15 different flow-channels with a snake-type layout and that gives a total of 122,880 trenches for one strain, which is crucial for detecting rare events like persister states. (d) The throughput of the experiment depends on imaging conditions required for that particular type of experiment. In a single-color fluorescent experiment using an air objective we can achieve the maximum throughput of tracking 131,072 cell lineages every 5 min. As the number of colors channels to be imaged increases, the throughput decreases, as the filter change and exposure times of each color contributes to the delay causing total number of cells imaged every 5 minutes to be lower. For phase contrast imaging, we are interested in tracking individual cell growth and division at least once every minute. So the throughput is lower (30,000 cells per 1 min using a 40x air phase). To get high-resolution images that enable accurate tracking and segmentation, we have used a high magnification (100x) oil lens. This further reduces the throughput and we can track 3,500 cells every one minute.

### Section 7 Comparison of bulk growth with growth in microfluidic devices

In order to monitor the bulk-growth of the culture over time, we have used a custom-designed optical density recording system (OD-meter). The OD-meter is designed as a housing for the bubble-trap described in Section 2. The detector—placed at 180° with respect to the emitter—measures the transmission of light through the sample, which serves as a metric for OD ($OD_{600}$) after calibration. In Figure S7a, we show example plots of $\log_2(OD_{600})$ over time for cultures growing in different media composition and temperatures. The growth-rate in a particular growth-condition (media/temperature) is estimated from the gradient of the steepest part of the $\log_2(OD_{600})$ curve. The corresponding single-cell estimates are calculated from single-cell growth rate obtained from length measurements of individual cells over time. The average growth-rates from the bulk measurements (x-axis) and the single-cell size measurements in the microfluidic device (y-axis) are compared in Figure S7b.

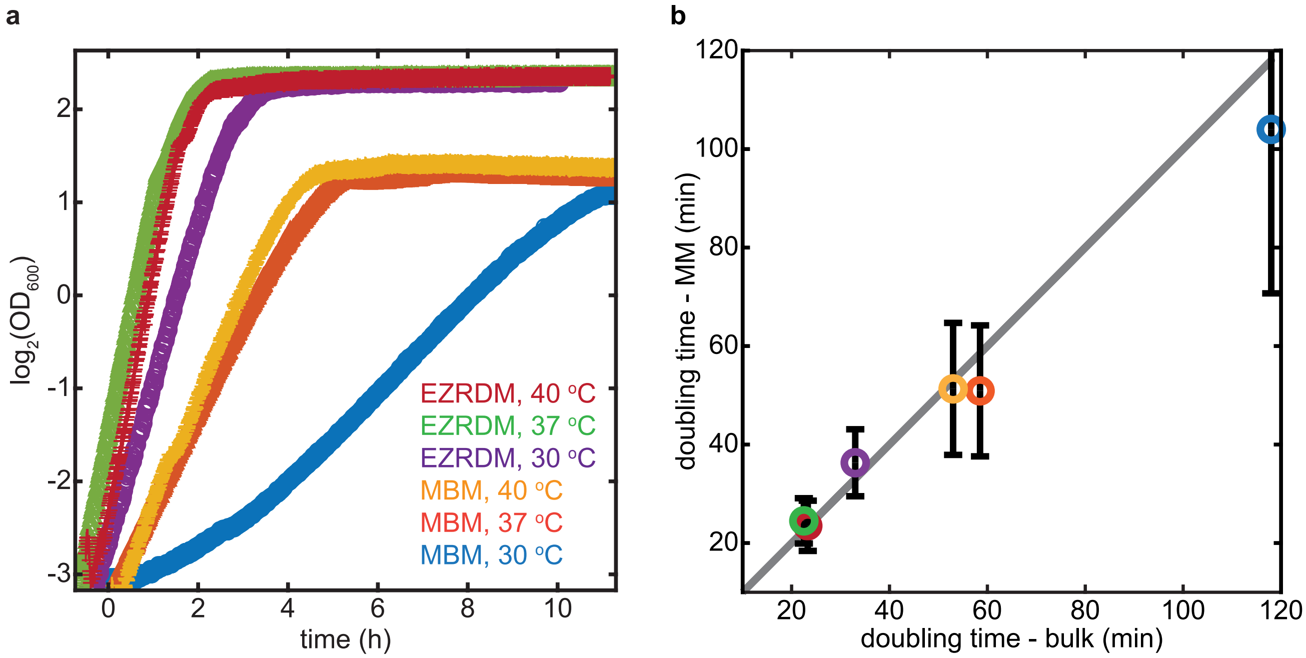


**Figure S7.** (a) Logarithm of $OD_{600}$ vs time for cells growing under six different growth-conditions (two media: rich-defined (EZRDM) and poor-defined (MBM) and three different temperatures: 30°C, 37°C, and 40°C). (b) Growth-rates from single-cell size measurements in mother machine (y-axis) are compared with bulk OD measurements (x-axis) for all six conditions. The grey-line corresponds to $y = x$.

### Section 8 Comparison of growth-dynamics in the bulk culture and at single-cell level

Here we compare the growth dynamics at single-cell level (measured from change in cell length over time) and at bulk level (measured by rate of change in $OD_{600}$ over time). In Figure S8a, we plot in blue the average length of cells loaded in the mother machine as they traverse the growth-curve and in black the $OD_{600}$ of the growing culture that flows through the microfluidic device. The two datasets are shifted by the delay time of ∼20 minutes (time for the culture to flow from the flask to the microfluidic device). However, it is important to note that the delay is likely to be shorter than the time it would need to flow from the flask to the mother machine, since the cells continue to grow in the tubing, which is maintained at same temperature (37°C) throughout the entire path. Until t = 8 h, cells are in exponential phase, and grow with an average size of 5.81 $\mu m$. After the diauxic shift (t = 9.1 h), the cell size starts to decline as the bulk growth also continues to decline. Near t = 11 h into the experiment, the bulk-growth stops (OD reaches plateau), but cell size continues to decrease for a while. Examination of the single-cell length traces suggests that the specific growth-rate ($g(t) = \mathrm{d}\log(t)/\mathrm{d}t$) of the cells also halts near that time, but some of the cells stochastically decides to divide one more round, resulting in smaller cells deeper into stationary phase (Figure S8b).

Such discrepancy between the growth and division is also seen during the diauxic shift. Until t = 9 h cells continue to grow and divide with a rate consistent with doubling the volume of a single cell and their number every ∼23 minutes. Right after the diauxic shift the specific growth-rate sharply declines across the population (Figure S8c). The sharp decline in specific growth is perfectly aligned with the drop in bulk growth measurements (grey-line), and always coincides with the diauxic shift observed in OD curves. In Figure S8d, we show two example single-cell length traces that demonstrate how growth and division becomes decoupled after the diauxic shift. Starting from t= 8.9 h until t =9.6 h, cells continue to divide every ∼23 minutes, while the specific growth remains slow. This phenomenon is reminiscent of rate-maintenance resulting from the processive nature of replication machineries [7]. The reduction of growth-rate and near constancy of splitting rate causes the cells to decrease their size rapidly. After this period, the specific growth-rate and the splitting rate both declines and cells halt growth and division. A fraction of cells stochastically divides to create two daughter cells of smaller, likely resulting from unfinished rounds of replication (Figure S8b).

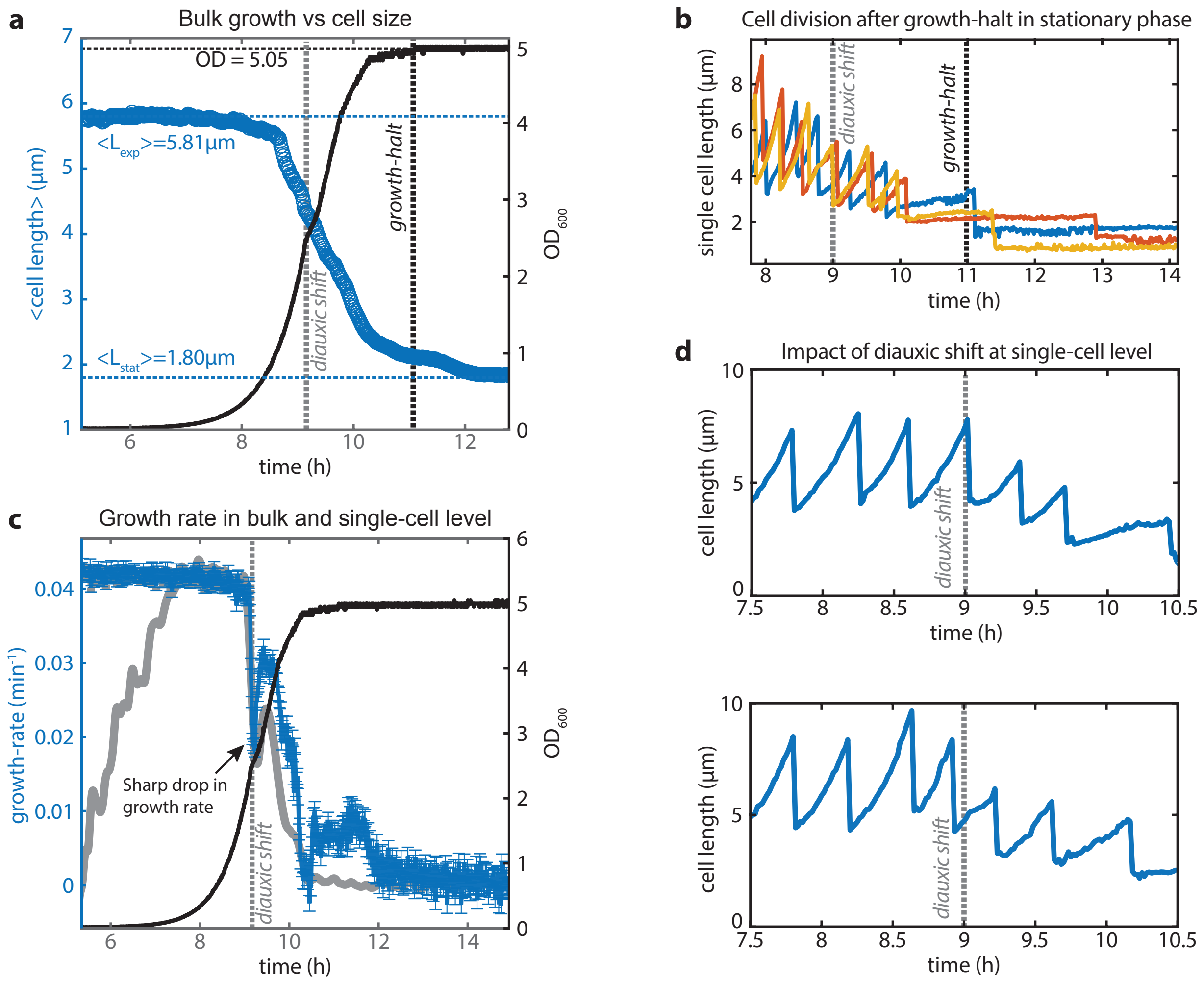


**Figure S8.** (a) Comparison of the dynamics of optical density of the bulk culture and average length of cells trapped in mother-machine during entry to stationary phase. The bulk culture passes the diauxic point slightly after t = 9h and then growth resumes for another 2 hours before the media is exhausted at t = 11 h and the bulk growth halts. The average cell length $\langle \mathrm{L(t)} \rangle$ continues to decline for a few hours after this due to stochastic division of cells in the absence of any growth, as shown in (b). (c) Plots of the average specific growth-rate of single cells ($\langle g(t) \rangle = \langle \mathrm{d}\log(L(t))/\,\mathrm{d}t \rangle$, average rate of change of log of cell length, in blue) over time with the bulk growth-rate ($\mathrm{d}\log(OD_{600}(t))/\,\mathrm{d}t$, rate of change of $\log(OD)$, in grey). The average growth-rate in bulk culture (0.042/minute) nicely matches the single-cell growth-rate in the mother-machine (0.043/minute) before the diauxic shift. Both sharply decline during the diauxic point, and resume after $\sim$40 minutes. (d) Sample single-cell length traces during showing the discrepancy between growth and division after diauxic shift. Cells slow down growth right after passing through diauxic shift, but keep dividing every $\sim$23 min, essentially behaving like a ‘timer’. The ‘timer’ mode lasts for two divisions, equivalent to the time needed to finish one round of replication.

### Section 9 *B. subtitlis* and *E. coli* both follow adder mode during exponential phase

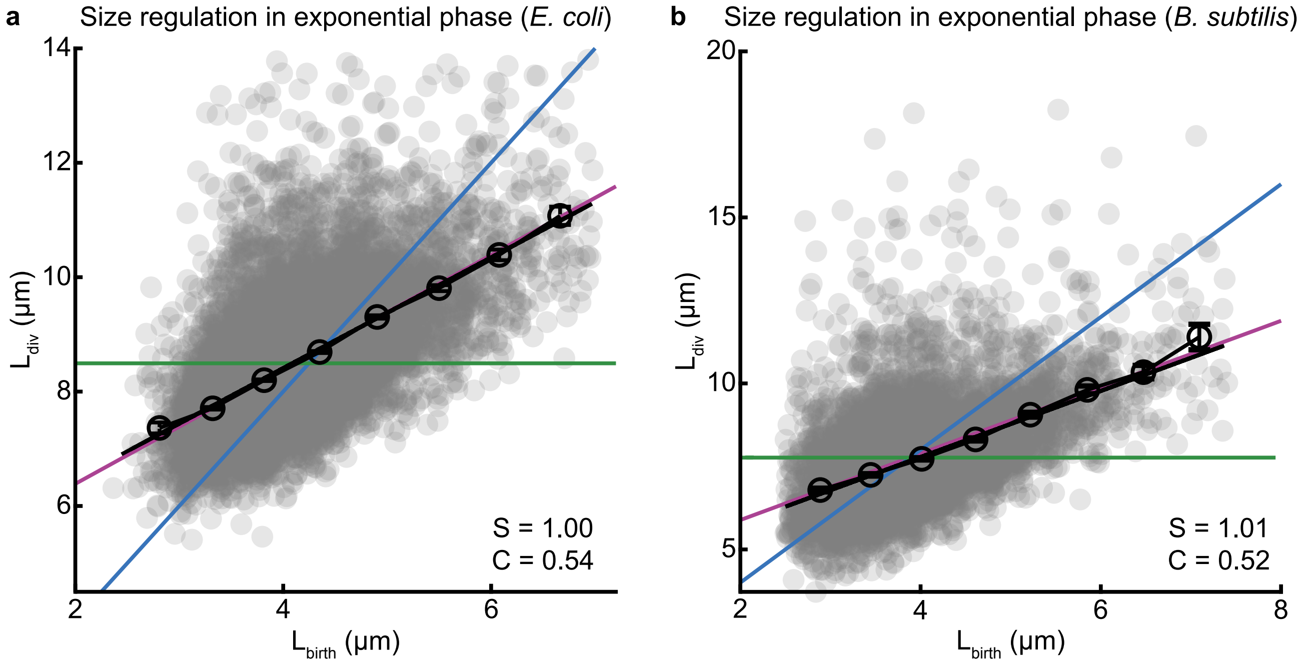


**Figure S9.** (a) ***E. coli.*** Correlation between cell lengths at birth and division are plotted during exponential growth phase. Grey circles are data from single division events, and black circles are binned along the x-axis. Theoretical lines for the classic modes - timer, sizer, and adder - are shown as blue, green, and purple lines. (b) ***B. subtilis.*** Correlation between cell lengths at birth and division are plotted during exponential growth phase.

### Section 10 Transition from adder-mode towards timer-mode during entry to stationary phase

The high throughput of our measurement platform allows us to bin the data in narrow time-windows to avoid averaging over continuously changing conditions. To investigate whether the cells are gradually or abruptly switching to the mixed timer-adder mode during the entry to stationary phase, we have binned the data in 2 min intervals. We have analyzed the correlation between the length at birth and at division for cells born within that interval. We see that after the diauxic shift ($t = 9h$) cells gradually switch from adder mode ($S = 1, C = 0.5$) towards the timer mode ($S = 2, C = 1$). This suggests that cells become less corrective as they enter stationary phase.

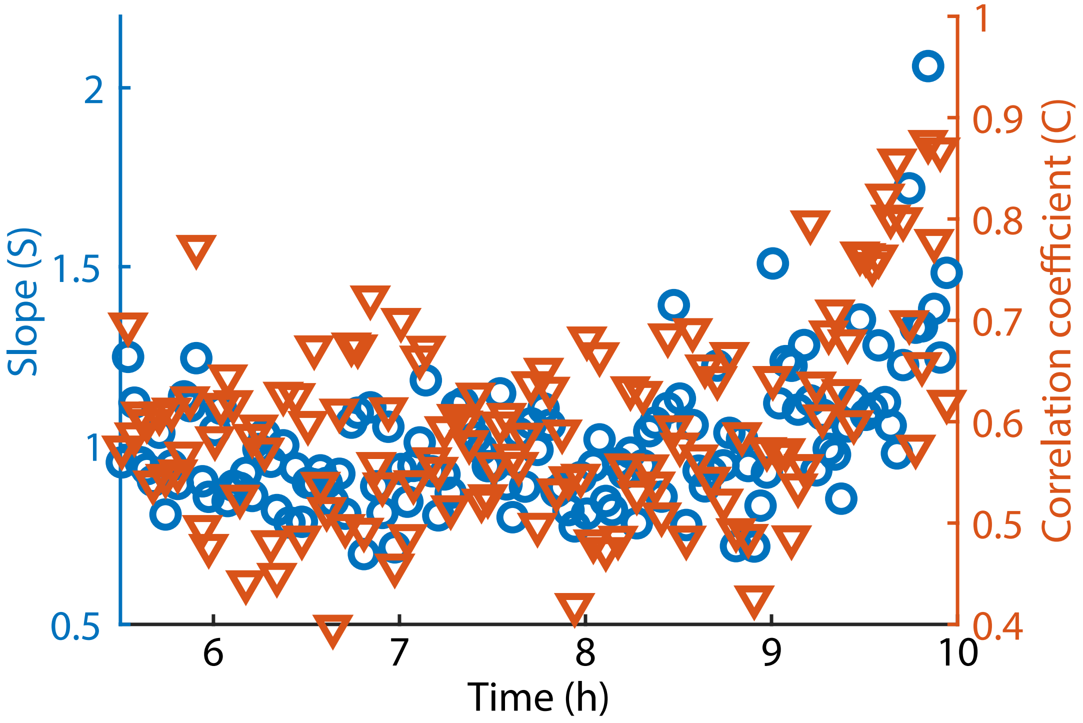


**Figure S10.** Division events are binned at 2 min intervals to examine the mode of size regulation at each interval. This avoids mixing cells from different part of the growth-curve where the averages can differ significantly within the data. The plot of the slope ($S$) of the trend-line and the correlation coefficient ($C$) between the size at birth and the size at division demonstrate gradual transition from adder mode in exponential phase (before $t = 9$ h) to a mix of timer and adder mode after cells pass through the diauxic point (after $t = 9$ h).

### Section 11 *B. subtitlis* behave as sizer during exit from stationary phase

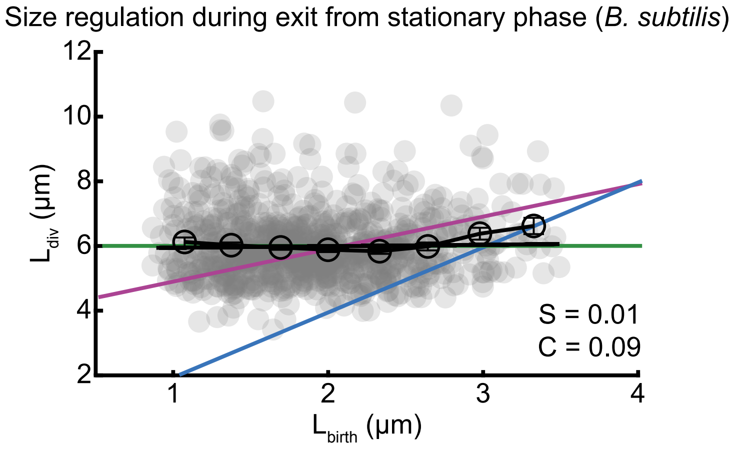


**Figure S11. *B. subtilis*.** Correlation between cell lengths at birth and division are plotted during the exit from stationary phase.

### Section 12 PSF artifacts in agar-pad images

To ensure that the cells in mother-machine are representative of cells in the batch culture, we collected cells from batch cultures and compared their size and gene-expression with cells growing in the mother machine at the same time points (see Figure 2b-d, main text). Except few points near the entry to the stationary phase, the cell size and gene-expression data from the mother-machine and the agar-pads matched very closely (Figure 2d). When we investigated the discrepancy near the stationary phase, we found that cells were present mostly in clumps. As mentioned earlier (Section 5), the packing of multiple cells close to each other causes light bleed-through from diffraction. The fluorescence intensity of a cell's immediate neighbors can therefore distort its measured intensity. Investigation of the cells on agarose pad at different densities revealed that the average intensity of densely packed cells was nearly twofold higher than that of sparsely distributed cells (Figure S12b). Therefore, we concluded the origin of the discrepancy was not due to the artefact of the mother-machine platform, but rather from point spread function in agar-pad imaging.

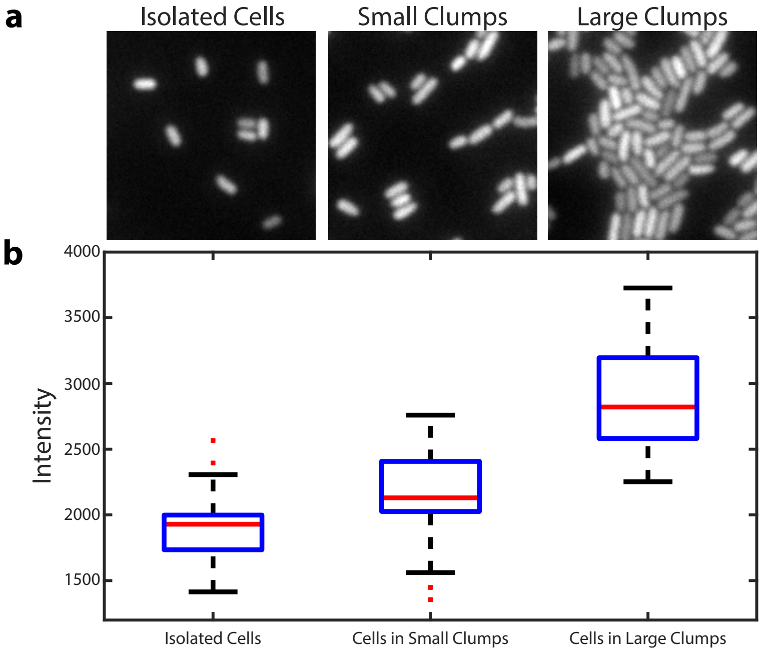


**Figure S12.** (a) Example images of cells on agar-pad. We chose three class of cells to examine the dependence of the average intensity from a cell on the size of clump it belongs to. Left: isolated cells, middle: cells in smaller clumps consisting of two to three cells, right: bigger clumps containing 20-50 cells. (b) Average intensities of the cells in each class is plotted. This shows that typical average from an agar-pad data set from cells in stationary phase will over-estimate the true average up to 1.5 fold, accounting for the discrepancy observed between the mother-machine and batch-culture data in Figure 2c, main text.

### Section 13 Data quality, analysis, and growth-measurements

For size regulation analysis, we have used a 100x phase contrast oil objective for imaging, which gives $0.065\mu m/$pixel size. The data were collected with a frame-rate of 1 minute/frame. In a typical experiment, we collect data for $\sim$3,500 single lineages with these settings. Roughly 50-60% of the cells stay in the trench for the entire duration of the experiment (36 h), while the rest get washed away due to filamentation or chaining. Raw data from a typical single-cell length trace is shown in Figure S13a. Segmentation error causes rare but prominent anomalies in the trend (see red arrows in (a)). In the pre-processing stage, we use a custom-build filter to remove such errors, before we perform the size-regulation analysis. A zoom of the trace in the stationary phase is shown to give a sense of the quality of size estimation from the segmentation protocol (b), since there is no growth during stationary phase ($\langle g(t)\rangle = 0.0001$/minute). We estimated the segmentation error from the standard deviation of cell-lengths from individual traces during stationary phase, and found it to be subpixel (average $\sigma = 0.7$ pixel) and independent of the cell length (c). Since the average length in exponential phase is near 80 pixels, the expected error is less than 1%. A zoom of the trace in the exponential phase is shown (d). We estimate the division times from the inter-peak distances and the distribution of all such events from a single experiment is shown in (f). The long tail is due to transient chaining events. The cell length increases exponentially with the time between two consecutive division events. A collection of such intervals from the trace in (a) is shown in (e). The specific growth-rate g(t) is estimated from the slope of the $\log_2$(cell-length) at each interval (f). The distribution of estimated growth-rates from all such intervals from one experiment is shown in (g). The mean growth-rate is 0.044/minute, which corresponds to a size doubling time of 22.7 minute and is remarkably consistent with the average number doubling time or division time $23.3 \pm 5.6$ min.

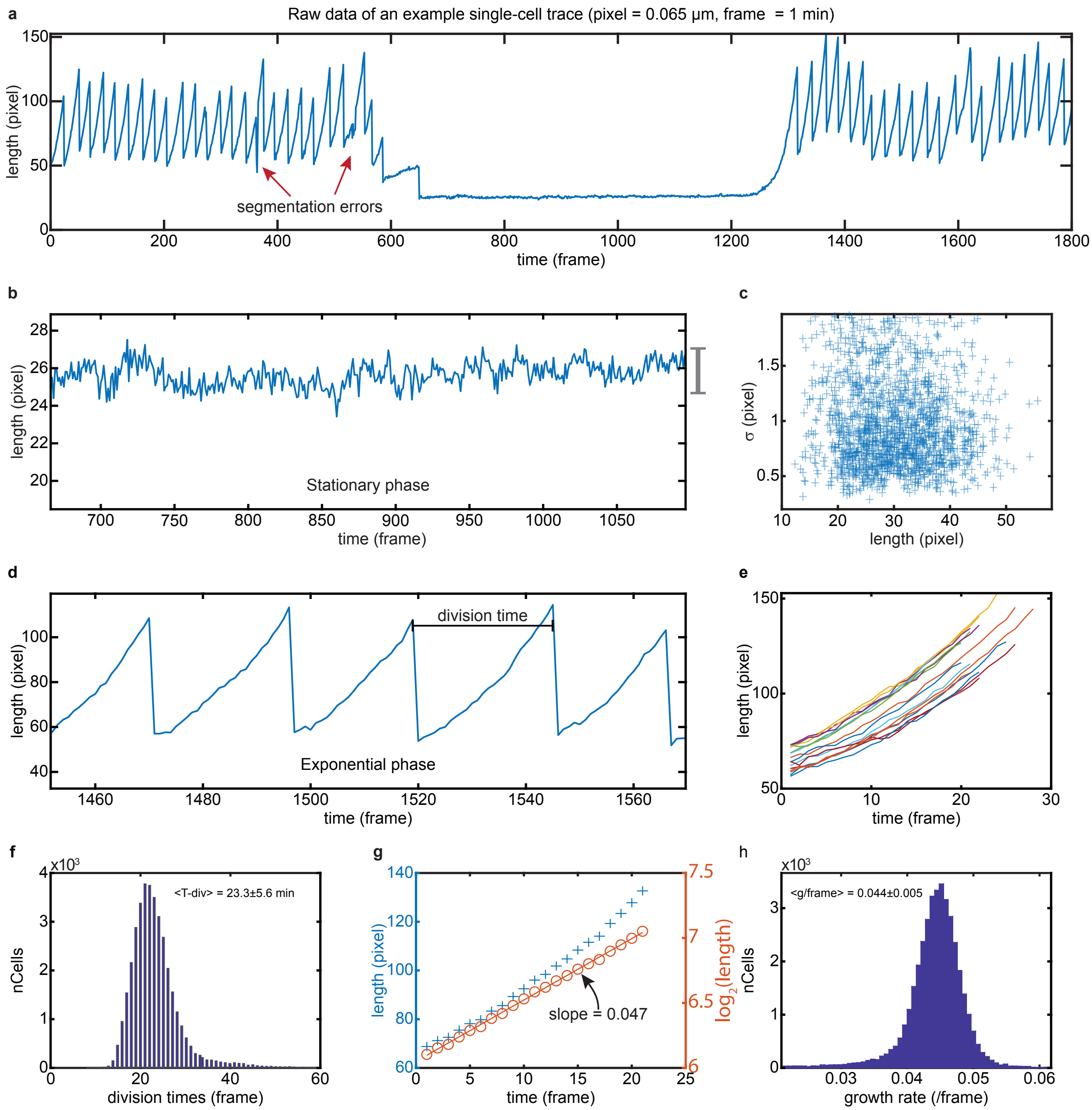


**Figure S13.** (a) Example: raw-data from a single-cell trace is shown during entry and exit from stationary phase. (b) Zoom of cell-length from the stationary phase is shown to illustrate the magnitude of fluctuations in the measurements. Measurement error is estimated from each trace as the standard deviation of lengths in stationary phase. In (c) we plot the measurement error vs average cell length for each trace in stationary phase. Cell-lengths and measurement errors are reported in pixels (1 pixel = 0.065 $\mu m$). (d) A zoom of the data from exponential phase is shown. The time-gap between two consecutive peaks is used as an estimate for division times. A collection of time-series of cell lengths within such intervals from a single-cell length trace is shown in (e). The length seems to increase exponentially over time. (f) Distribution of all the division times estimated from all the cells (N = 2,215 ) imaged in this experiment. (g) The specific growth rate for each cell during a particular interval is calculated from the slope of the $\log_2$(cell-length) vs time plot. The raw cell length is plotted in left-axis (blue crosses), while $\log_2$(cell-length) is plotted in the right-axis (orange circles). The slope for this particular trace is 0.047. (h) Distribution of slopes (specific growth-rate) from all such intervals from all the cells in this experiment is shown.

### Section 14 Rate-maintenance and cell size in stationary phase

The 'rate-maintenance' phenomenon causes cells to divide and become smaller during the entry to stationary phase. However, since this is a consequence of multifork replication, which in turn is consequence of fast growth in rich conditions, we examined if the size-reduction during entry to stationary phase ceases in poor growth-conditions. In Figure S14, we plot the distribution of cell sizes in different growth-phases and during birth-division for two different media conditions. The summary statistics of the distributions are provided below (Table S1). To quantify the net size reduction resulting from rate-maintenance we compare the distribution of birth-sizes to the average sizes in stationary phase, since cells don't grow and divide in the stationary phase. In case of rich defined medium (EZRDM), cells become 2.2x smaller in stationary phase compared to the average birth-size in exponential phase. This size-reduction factor was same for all three different temperatures examined in this paper (30, 37 and 40°C). However, in poor defined medium (MBM), the size distribution of stationary phase and birth-size in exponential phase match closely, supporting the hypothesis that in the absence of rate-maintenance cells simply halt growth upon starvation. This suggests that at least under the conditions examined in this paper, rate-maintenance is the major if not the only mechanism responsible for cell-size drop during entry to stationary phase.

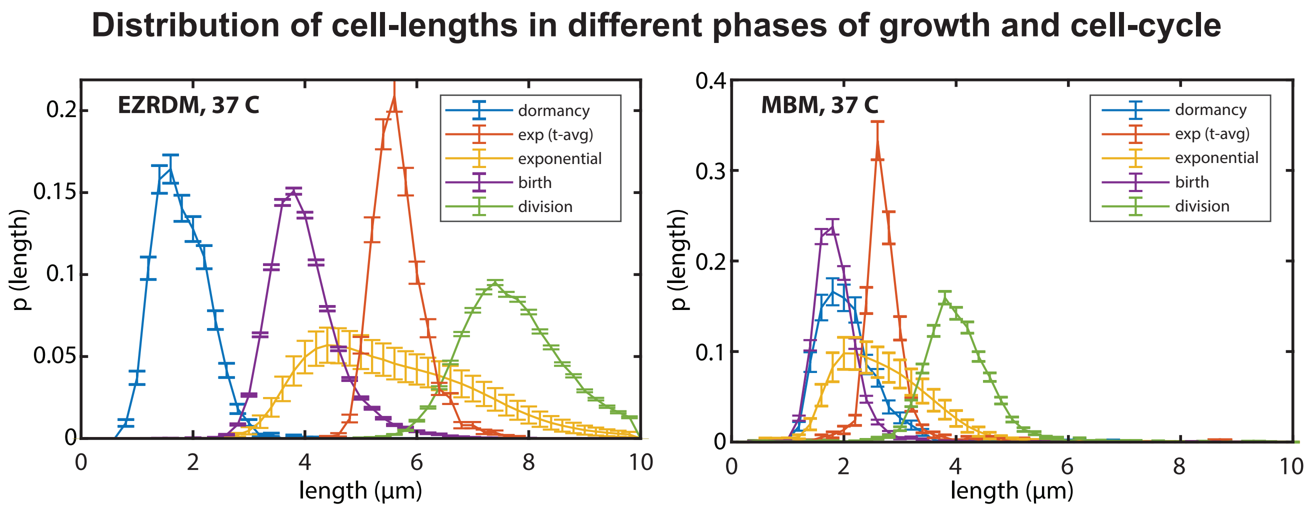


**Figure S14.** Comparison of cell size distribution in different phases of growth-curve (during dormancy or stationary phase, and in exponential phase), and birth-division sizes in exponential phase. The observed broad distribution of cell-size in exponential phase is a result of the spread in size from birth to division, and a time-averaged value for each cell (orange curve) follows a much narrower distribution. Left: results relative to rich defined medium (EZRDM). Right: results relative to poor defined medium (MBM).

| Growth condition | $L_{birth}$ ($\mu$m) | $L_{div}$ ($\mu$m) | $L_{stationary}$ ($\mu$m) | $L_{exponential}$ ($\mu$m) | Doubling time (min) |
|---|---|---|---|---|---|
| EZRDM, 37°C | $4.05 \pm 0.61$ | $7.99 \pm 1.051$ | $1.90 \pm 0.51$ | $5.85 \pm 0.49$ | $23.5 \pm 5.1$ |
| MBM, 37°C | $1.99 \pm 0.82$ | $4.08 \pm 0.99$ | $2.08 \pm 0.40$ | $2.78 \pm 0.29$ | $50.9 \pm 13.3$ |

**Table S1.** Statistics of distributions in rich defined medium (EZRDM) and poor defined medium (MBM).

### Section 15 Size regulation in different temperatures

| Growth condition | Doubling time (min) | C (exponential | C (entry | C (exit |
|---|---|---|---|---|
| EZRDM, 37°C | $23.5 \pm 5.0$ | 0.51 | 0.74 | 0.10 |
| EZRDM, 30°C | $36.3 \pm 6.8$ | 0.45 | 0.55 | 0.11 |
| EZRDM, 40°C | $24.5 \pm 4.6$ | 0.50 | 0.67 | 0.04 |

**Table S2.** Comparing cells in rich defined medium (EZRDM) at three temperatures shows that the size-regulation in different growth-phases does not change significantly with temperature. According to the 'rate-maintenance' mechanism, we expect the size-regulation during entry to stationary phase to depend on the replication frequency. Consistent with this, for T = 30°C, where the doubling time is comparable to the replication time, the 'timer'-like aspect during entry almost disappears.

### Section 16 Bulk measurements of persisters

To estimate the persister frequencies in bulk cultures, we followed the protocol described by Balaban *et al.* [8]. A flask with 50 ml of fresh media, either Luria-Bertani (LB) or EZRDM (EZ Rich Defined Medium) supplemented with F108 (0.8 mg/l) was inoculated with 10 $\mu m$ of an overnight culture of the SB7 strain (please see Section 18 for details). Cultures were incubated at 37°C and 220 rpm for 24h or 48h. Once the incubation period was over, 50 $\mu m$ of the stationary-phase culture was diluted into 5ml of the corresponding fresh media. At this point a sample of 100 $\mu m$ was taken, serially diluted and plated to determine the number of colony forming units (cfu) at $t = 0.5\mu m$ of ampicillin (50 mg/ml) was then added to each tube containing the 5 ml culture, and incubation was resumed for 3 hours, after which a 100 $\mu m$ sample was taken, serially diluted and plated to determine the cfus of persisters. The results are shown in Table S3.

| **Growth medium** | **Stationary phase duration = 12h** | **Stationary phase duration = 36h** |
|---|---|---|
| LB | $\mu = 1.7 \cdot 10^{-4}$; $\sigma = 0.5 \cdot 10^{-4}$ | $\mu = 3.7 \cdot 10^{-3}$; $\sigma = 0.6 \cdot 10^{-3}$ |
| EZRDM | $\mu = 2.5 \cdot 10^{-4}$; $\sigma = 1.0 \cdot 10^{-4}$ | $\mu = 9.2 \cdot 10^{-4}$; $\sigma = 0.6 \cdot 10^{-4}$ |

**Table S3.** Mean ($\mu$) and standard deviation ($\sigma$) of estimated frequency of persisters.

### Section 17 Growth-curve experiments *B. subtilis*

The growth-curve experiments of *B. subtilis* were done with similar microscopic settings as the ones performed with *E. coli*. The main differences were in the (a) strains used for the work, (b) protocols for growing the culture and (c) the microfluidic devices used for loading the strains that were imaged along the growth-curve. We give details of each of these three aspects below.

*Strains:* We repeated the bacillus experiments with strains from two different backgrounds (3610 and 168) and obtained similar results. The work presented in this paper are using strains from 168 background. We used the wild-type 168 strain in the culture flask, and a strain BGD12 in the mother machine. The strain BGD12 contains a constitutive mCherry expression cassette, and a straight flagellum mutation (hagA233V) that prevents the flagellum from generating force so that motile cells cannot swim out of the microfluidic channels [5].

*Growth-curve:* Cells were grown in a rich defined medium that was optimized for *B. subtilis* (Lab protocol). The average division time of *B. subtilis* cells in this medium at 37°C in our setup is 21 min. We added 0.1% BSA in the culture flask to prevent cells from sticking to the tubing in the flow path9. Since *B. subtilis* cells require more aeration than *E. coli* cells, the volume of culture in the flask was reduced (100ml culture in 500ml baffled bottom flask) to increase surface-to-volume ratio.

*Sample preparation:* Since *B. subtilis* cells are slightly longer and wider compared to *E. coli* cells in the exponential phase, and tends to chain more frequently, we used a different microfluidic device design for the experiments with *B. subtilis* cells. For the experiments described in this paper, we chose trench dimensions that are optimal for the dimensions of *B. subtilis* cell in exponential phase, where cells are largest, and to minimize loss events from stochastic chaining (trench-length = 35 $\mu m$, trench width = 1.7 $\mu m$, trench height = 1.3 $\mu m$). The surface of the microfluidic devices were passivated with 0.1% BSA, as described in our previous work [5]. Chips were spun at 5000 rcf for 10 min to load the cells and cells from the feeding channel were washed with fast flow of media from a syringe (40 $\mu m$/min) and then the flow-rate was reduced to 20 $\mu m$/min for the rest of the experiment.

### Section 18 Strain construction control

For all the *E. coli* experiments described in this paper we have used the MG1655 7740 background, which was used by the Blattner lab for the complete sequencing of *E. coli* K-12. However, this strain has an insertion of IS1H at flhDC, which causes hyper-motility and therefore causes the bacterial cells to swim out of the mother machine trenches. To avoid this, we have inserted ΔmotA using P1 transduction from the corresponding strain in keio collection [9]. The strain SB8 used in all the size-regulation work described in this work is MG1655 7740 with ΔmotA. The size-regulation work involved only phase contrast microscopy, and therefore did not require any fluorescent markers. For the stress-response work, we used a strain that constitutively expresses mCherry1-11-mKate from the glmS site. This strain (SB7), constructed by integrating glmS::PRNAI-mCherry1-11-mKate-T1 terminator-FRT Kan FRT::pstS into the SB8, enables us to get a bright fluorescence image of cell in RFP channel and is used for single-cell segmentation. To ensure the measurements performed in this study did not have artefacts from the ΔmotA or the mCherry1-11-mKate expression, we have performed control experiments to check and validate the results with other strains. We have compared the results of stress-response dynamics along the growth curve between ΔmotA and ΔfliC, which also renders the cell non-motile, and finally compared with the MG1655 6300, which is much less motile than 7740 version and found that the results were identical. We have also checked the frequency of persisters between the wild-type cell (MG1655 6300) and the strain used in the persister experiments (SB7) using bulk experiments and found the frequencies to be same (0.01 for both after overnight stationary phase and 3 hours long treatment with 50 $\mu m$/ml ampicillin).

### Section 19 Media Recipes

**EZRDM (EZ-Rich Defined Medium, optimized recipe).** The medium is based on the recipe developed by Neidhardt *et al.* [10]. The MOPS EZ Rich Defined Medium Kit is available commercially (Teknova, #M2105) and was optimized for osmolarity with 5M NaCl according to work by Konopka *et al.* [11]. For 1L of media mix together the following components:

| | |
|---|---|
| 10X MOPS | 100ml |
| 5X EZ Supplement | 200ml |
| 10X ACGU | 100ml |
| 0.132 $MK_2HPO_4$ | 10ml |
| 20% Glucose | 10ml |
| 5M NaCl | 15.2ml |
| Sterile $H_2O$ | 564.8ml |

Filter the medium using sterile filters (0.22 $\mu m$) and store in smaller batches at 4°C (up to a week) or at -20°C (up to few weeks).

**MBM (Poor Defined Medium, optimized recipe).** The medium is based on the recipe developed by Neidhardt *et al.* [10]. The MOPS Minimal Defined Medium Kit is available commercially (Teknova, #M2106) and was optimized for osmolarity. For 1L of media mix together the following components:

| | |
|---|---|
| 10X MOPS | 100ml |
| 0.132 $MK_2HPO_4$ | 10ml |
| 20% Glucose | 10ml |
| 5M NaCl | 19.6ml |
| Sterile $H_2O$ | 860.4ml |

Filter the medium using sterile filters (0.22 $\mu m$) and store in smaller batches at 4°C (up to a week) or at -20°C (up to few weeks).

**Pluronic Solution.** Mix 100gr of Pluronic F108 (Sigma Aldrich, 542342-250G) in 1L of sterile $H_2O$ and mix with a magnetic stir bar. Once dissolved, filter the solution using 0.22 $\mu m$ sterile filters and store at room temperature.
